# Systems biology and machine learning approaches identify metabolites that influence dietary lifespan and healthspan responses across flies and humans

**DOI:** 10.1101/2023.07.09.548232

**Authors:** Tyler A.U. Hilsabeck, Vikram P. Narayan, Kenneth A. Wilson, Enrique Carrera, Daniel Raftery, Daniel Promislow, Rachel B. Brem, Judith Campisi, Pankaj Kapahi

**Affiliations:** Buck Institute for Research on Aging, Novato, California, 94945, United States; Davis School of Gerontology, University of Southern California, University Park, Los Angeles, California, 90089, United States; Dominican University of California, San Rafael, California, 94901, United States; Northwest Metabolomics Research Center, Department of Anesthesiology and Pain Medicine, University of Washington, Seattle, Washington, United States; Department of Pathology, University of Washington, Seattle, WA 98195, United States; Department of Biology, University of Washington, Seattle, WA 98195, United States; Department of Plant and Microbial Biology, University of California, Berkeley, Berkeley, California, 94720, United States

**Keywords:** aging, dietary restriction, lifespan, metabolome, healthspan, Drosophila, GWAS, Mendelian randomization, machine learning, random forest modeling

## Abstract

Dietary restriction (DR) is a potent method to enhance lifespan and healthspan, but individual responses are influenced by genetic variations. Understanding how metabolism-related genetic differences impact longevity and healthspan are unclear. To investigate this, we used metabolites as markers to reveal how different genotypes respond to diet to influence longevity and healthspan traits. We analyzed data from *Drosophila* Genetic Reference Panel strains raised under AL and DR conditions, combining metabolomic, phenotypic, and genome-wide information. Employing two computational methods across species—random forest modeling within the DGRP and Mendelian randomization in the UK Biobank—we pinpointed key traits with cross-species relevance that influence lifespan and healthspan. Notably, orotate was linked to parental age at death in humans and counteracted DR effects in flies, while threonine extended lifespan, in a strain- and sex-specific manner. Thus, utilizing natural genetic variation data from flies and humans, we employed a systems biology approach to elucidate potential therapeutic pathways and metabolomic targets for diet-dependent changes in lifespan and healthspan.

---

Aging is the leading cause of morbidity and mortality in most developed countries. Dietary restriction (DR), restricting specific nutrients or total calories without causing malnutrition, has been demonstrated as a robustly conserved means to extend lifespan and healthspan. Diet-responsive pathways such as the Target of Rapamycin and Insulin-like signaling have been implicated as driving factors of lifespan extension by DR; however, variation in response to diet between different individuals within the same species indicate that additional mechanisms influence diet response and remain yet to be elucidated^1^. Recently, genetic variation has been implicated as a significant driver of these phenotypic responses to DR. Studies in invertebrates^2^ and rodents^3^ have utilized systems biology approaches to identify additional DR-responsive factors that influence metabolic health and lifespan. Additionally, multi-omics approaches have identified novel metabolites that are regulated by genetic variation to influence longevity under DR^4^. Nevertheless, in 2020, Jin and colleagues highlighted the challenge of establishing a correlation between lifespan and metabolite traits. This difficulty arises from the strong association between the dietary restriction (DR) response in mean lifespan and the metabolite traits observed in flies following the standard diet (AL). To avoid the confounding effects of an AL diet on the DR response, Jin, *et al.* 2020 used the residuals from a simple regression to identify connections between metabolites and lifespan^4^. Meanwhile, another study conducted by Wilson, *et al.* (2020) highlighted a distinct absence of correlation between healthspan traits and lifespan traits, revealing disparate genetic regulators for each facet^2^. Despite the findings from these studies, it remains unclear how different phenotypic interactions occurring within an individual influence the relationship between health and longevity.

A powerful approach to exploring the relationship between health and longevity is to incorporate convergent data from multiple species. Research conducted using model organisms has identified many candidate genes and pathways responsible for the aging process^5,6^. Some of these genes and processes belong to evolutionarily conserved nutrient-sensing pathways, which suggests they may be relevant in humans as well. However, several large GWAS of human longevity show a surprising lack of association with lifespan-extending genes identified in model organisms^7^. The disconnect might in part arise from the effects of natural variation in a lifespan-associated gene, highlighting the importance of studying natural variation in model systems. An effective approach to investigate this variation across multiple species involves utilizing the metabolome as indicators of the pathways utilized in an individual’s dietary response, akin to “footprints”. While utilizing the metabolome has previously been done, as mentioned above, a fresh approach was required to establish a conserved connection between variations in metabolites with healthspan and lifespan.

Recent advancements in analytical techniques have enabled more comprehensive investigations into intricate traits. Machine learning, which leverages well-established high-dimensional datasets to create more accurate predictive models for specific phenotypes, has proven valuable in discovering novel factors impacting a range of biological traits, including those related to longevity and health^8^. However, it remains to be seen how the combined utilization of metabolomic and phenotypic indicators can transcend species and genotypes in response to dietary restriction (DR), enabling the prediction of specific elements contributing to various health and longevity attributes.

In previous work, we utilized the Drosophila Genetic Reference Panel (DGRP) to explore the effects of DR on metabolic phenotypes^9^, metabolome^4^, and lifespan and healthspan^2^. Across these datasets, we identified that no single metric could reliably predict lifespan, suggesting that multiple factors collectively influence the longevity outcomes of DR. Here, we use this data to probe the interplay between metabolomic, metabolic, and healthspan-related traits and lifespan. Employing random forest modeling, we pinpoint the relevance of metabolites orotate, threonine, and choline in constructing multiple lifespan models. This modeling approach circumvents potential issues arising from standard diet (AL) traits influencing DR response due to its bagging methodology^10–12^. Additionally, we uncover genetic candidates that may impact these associations, including *Src64B*, which demonstrates an association with mean lifespan. This study represents a pioneering effort, being the first to leverage genetic data from 174 strains, encompassing 34 distinct phenotypes and 111 measured metabolites within those strains, while also investigating targets within a human dataset.

To further validate the therapeutic potential of machine learning targets in humans, we performed Mendelian randomization using metabolomic data from 1,960 individuals in the UK Biobank. We evaluated the hypothesis that the metabolites identified through a random forest, together with their candidate genes influencing longevity in *Drosophila*, also exhibit associations with human lifespan. Collectively, these findings shed light on the intricate nature of dietary responses and their modulation by genetic variations. They specify the factors contributing to lifespan determination under DR (**Figure 1A**). This approach has led us to uncover pathways that could serve as valuable biomarkers of health and potential therapeutic targets for augmenting both human lifespan and healthspan.

**Figure 1.**
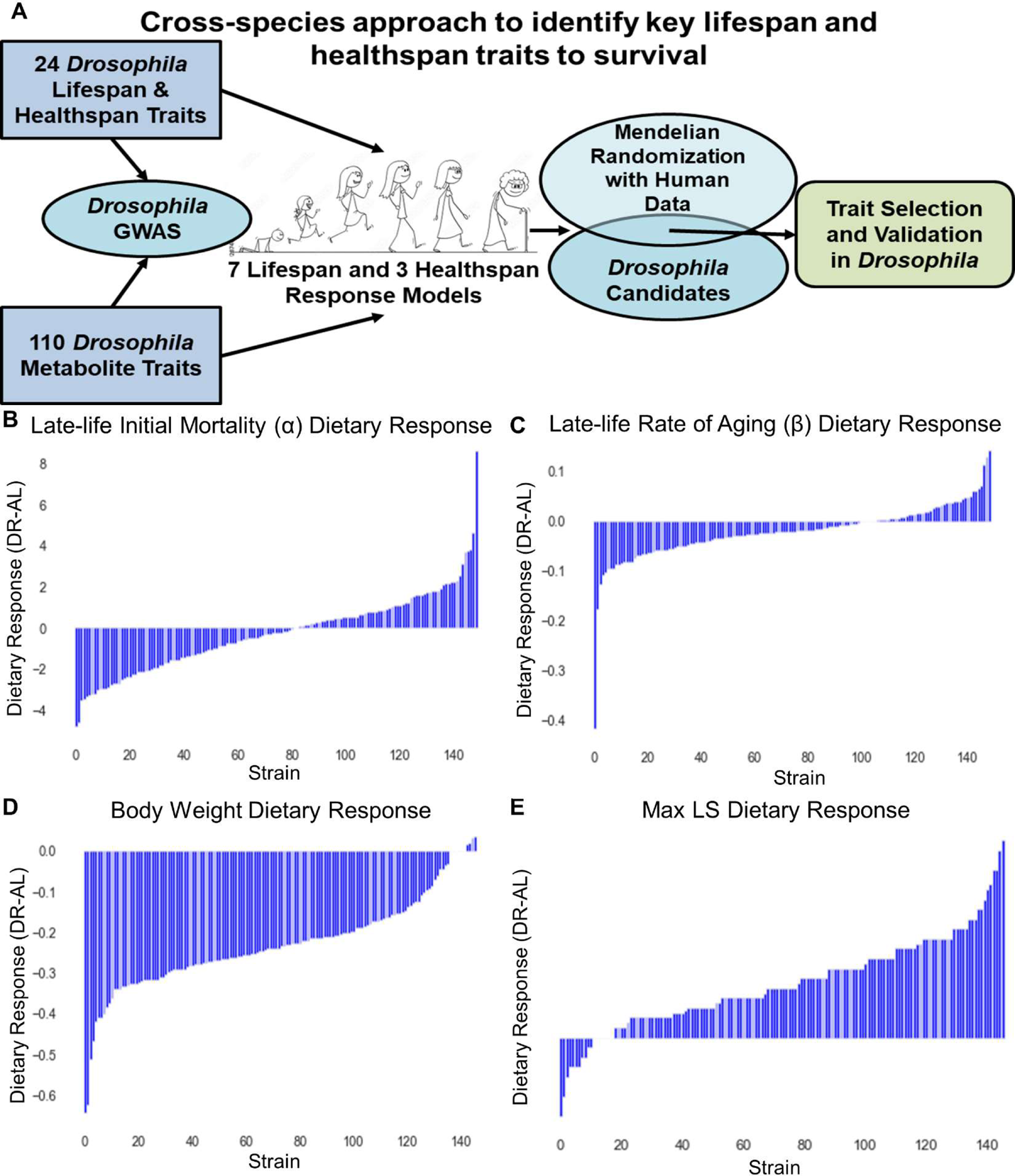
Diet influences Lifespan and Healthspan in a genotype-specific manner. DGRP show strong strain-specific responses to diet (A) Graphical workflow for modeling data from *Drosophila* and filtering traits using Mendelian randomization of human data. (B) DGRP strain traits on a high yeast (AL, red) or low yeast (DR, blue) diet were used to determine strain-specific dietary response (DR-AL). (B-E) Bar graphs of DGRP strain dietary response in (B) late-life initial mortality α, (C) late-life rate of aging β, (D) body weight (mg), and (E) max lifespan.

## RESULTS

### *DGRP* strains exhibit a wide variation in metabolite and phenotype responses to diet

We combined and reanalyzed data from previously published DGRP metabolite and phenotype datasets, including Nelson *et al*. 2016^9^, Jin *et al*. 2020^4^, and Wilson *et al*. 2020^2^, with flies fed both *ad libitum* (AL, 5.0% yeast extract) and DR (0.5% yeast extract) dietary conditions. From these data, we separated ten primary traits generally used in lifespan and healthspan determination studies as ‘response’ variables for modeling and used the remaining traits as ‘predictors’ (**Supplementary Table 1**). Principal component analysis (PCA) of predictors, after removing strains with incomplete data across all traits, failed to cluster into groups by diet. To determine if trait values from these strains would allow the data to cluster by diet, we imputed averaged values across all strains for these missing values (**Supplementary Tables 1&2**). Strains with missing values included four that lacked data for many phenotypes but had data for all metabolites and 33 additional strains with all metabolite data but missing at least one phenotype value (**Supplementary Table 1&2**). PCA of this imputed list clustered by diet and was used for downstream analysis, similar to what was previously shown^4^. Diet was part of principal component 1 (PC1), which explained ∼23% of the data’s variance. Approximately 32 principal components were sufficient to explain 85% of the variance in the data. Since diet explained a large portion of the variance, we determined the dietary response of each DGRP strain for each trait (value on AL subtracted from the value on DR, “DR-AL”). In general, DGRP strains responded to a DR diet similar to what has been seen before, with over 93% of strains having decreased body weight and at least 78% having an increase in most lifespan response metrics (**Figure 1B-E** and **Supplementary Figure 1**). These data demonstrate the variation in dietary response across the DGRP, and the few strains that do not have decreased body weight or increased lifespan from DR can give insight into the mechanisms underlying these and other positive effects of dietary intervention.

### Correlation of dietary response in DGRP strains identifies few metabolites that correlate with lifespan- and healthspan-related phenotypes

We looked for correlations between our predictor traits in each diet and the dietary response (AL, DR, and DR-AL), and found few correlations across data sets, with the majority of these being weak correlations (DR-AL, **Figure 2**). Just as observed by Jin, *et al.* in 2020, we also noted a moderate correlation (*r* = 0.38, P = 2.14 x 10^-6^) in median lifespan when comparing our “DR-AL” and AL datasets, and similar low to high correlations with the other lifespan metrics. We performed hierarchical clustering using Ward’s minimum variance method to form trait clusters. The strongest correlations on either diet alone (DR or AL) or used in the same data set (DR&AL), were between metabolites in the same pathway or between similar phenotypes (**Supplementary Figure 2A-C**).

**Figure 2.**
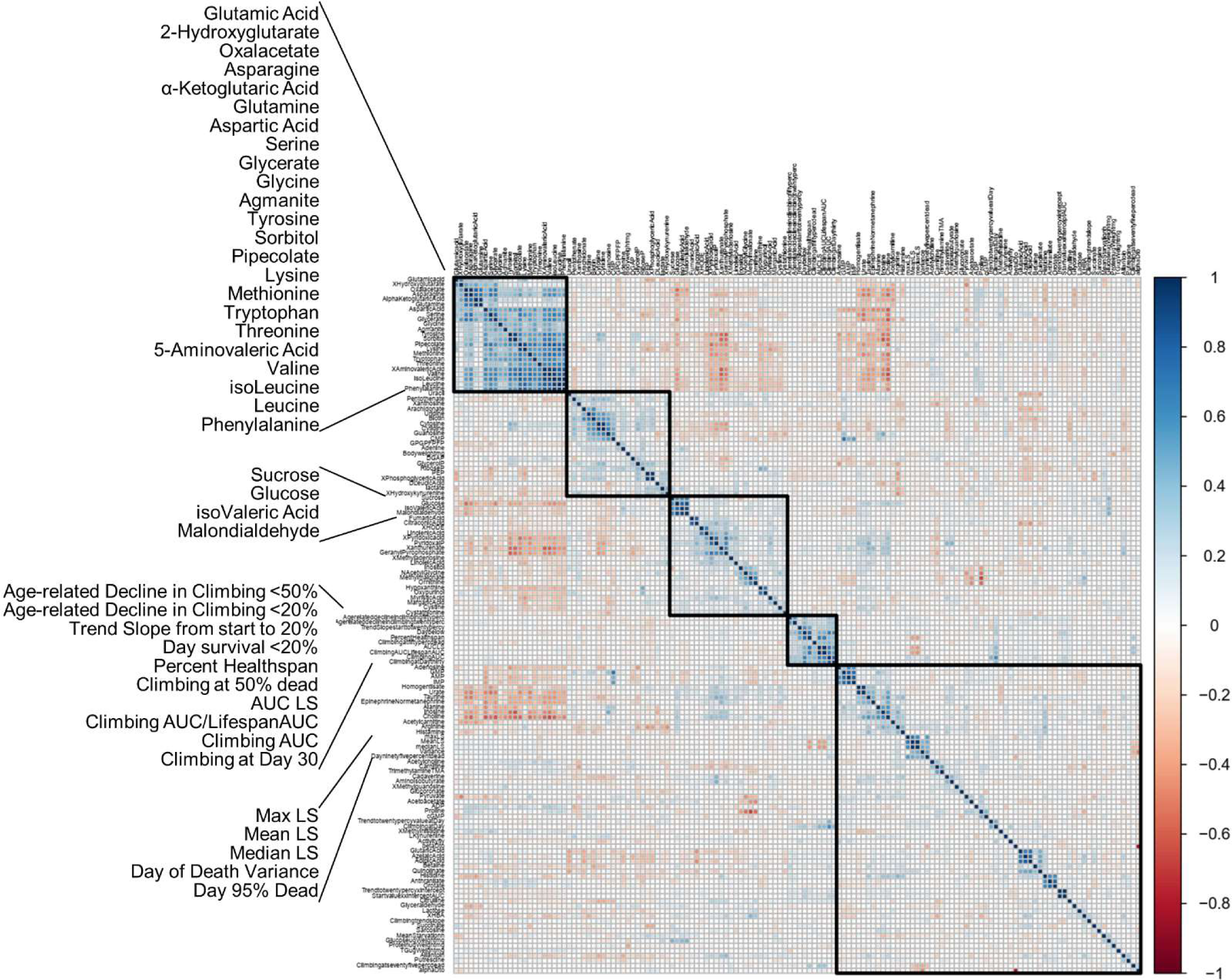
DGRP trait dietary responses correlate with similar traits, but few metabolites correlate with phenotypes. Heatmap of metabolite and phenotype dietary response correlations. Traits are clustered via hierarchical clustering using the Ward method, with 5 clusters highlighted on the diagonal. Names of similar traits within the same datatype that were correlated are highlighted as popouts. Pearson correlation values are shown as a gradient from 1 (dark blue) to -1 (dark red).

This trend continued for correlations in dietary response (DR-AL), though the top three correlations were between similar phenotypes rather than between metabolites. Unsurprisingly, the same pairs were strongly correlated in all comparisons. The lack of strong correlations between any of the predictor traits and the response traits points to a need for a different method for identifying factors that could predict and/or “build” these response traits.

### Random Forest modeling identified metabolite and phenotype traits used to build multiple lifespan response traits

To determine the predictor traits that contribute to determining the lifespan and healthspan response traits, we built random forest models for each response using all predictor traits as inputs. Predictors were used to build 10,000 initial models per response trait, with a final model being made based on the initial 10,000. Predictors used to build the final models were given an importance score based on the proportion of initial trees/estimators for which they were included. Models were created for each diet alone, DR & AL together, and the DR-AL response, and the important predictor traits were plotted for each response trait (**Figure 3, Supplementary Table 3**). Models for lifespan metrics in any dietary condition tended to be built with climbing-related traits, except the DR-AL response lifespan models which were also built by metabolite traits (**Figure 3A-F**). Specifically, threonine was used in at least 1% of the trees used for all 7 lifespan response models, peaking at 3% and 4% of the trees in the max lifespan and day 95% of flies dead models, respectively. Arginine was used in all lifespan models except for max lifespan, being in 4% of the late-life α trees, 5% of the late-life β, 4% of the day 95% of flies dead, and 3% of the variance in the day of death model trees. Choline was in 1% of late-life β, variance, and max lifespan model trees. Orotate was in 2% and 5% of late-life α and late-life β trees, respectively. Metabolites quinolinate, methylhistidine, and glyceraldehyde were also used to build at least 1% of trees for one or multiple lifespan models. The more traditional physical response traits (glucose levels, triglyceride levels, and body weight), were built predominantly by metabolites and protein levels (**Figure 3G-I**). Particularly, the models for glucose levels that included the AL diet had malondialdehyde as the most important trait.

**Figure 3.**
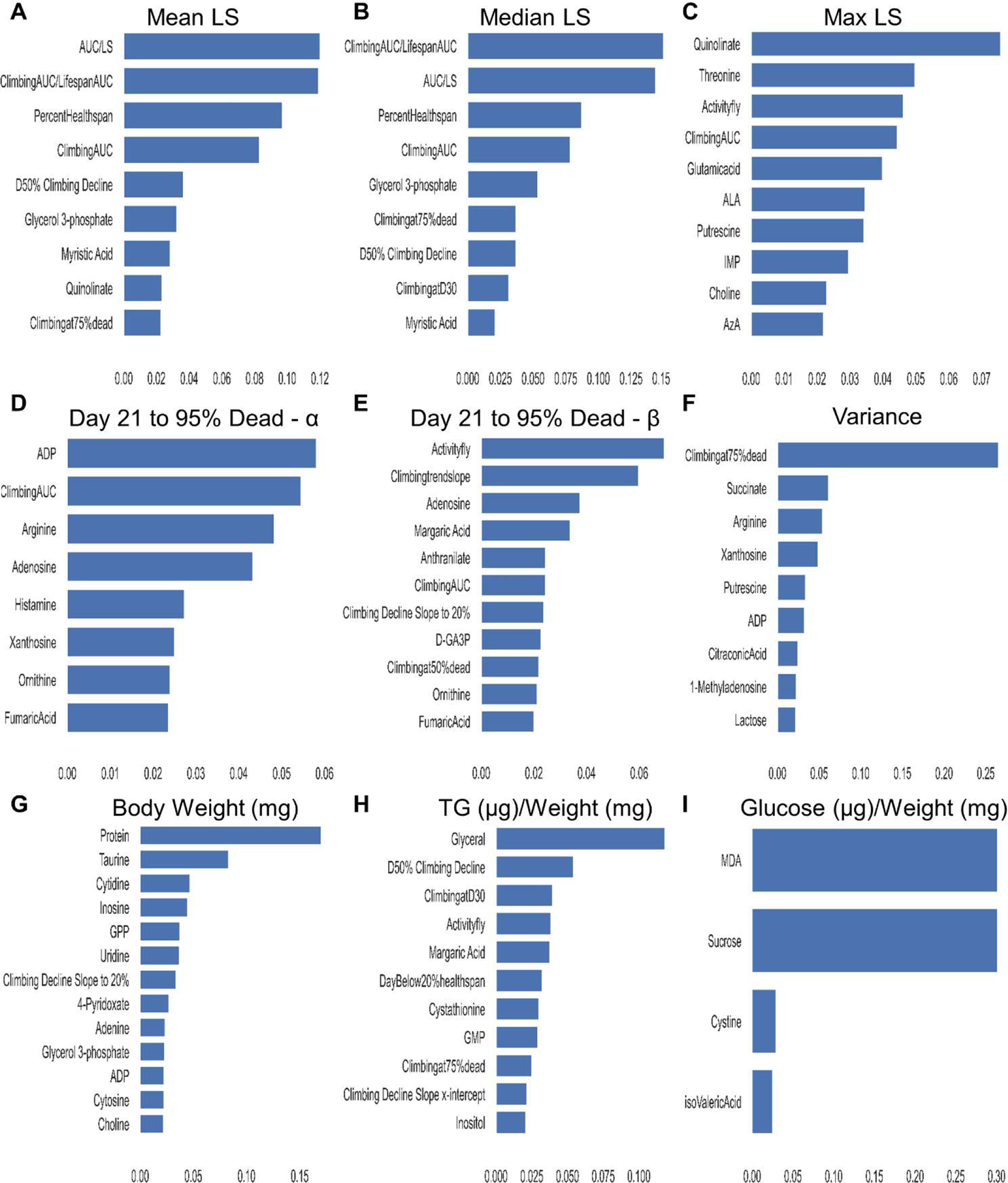
Random Forest models utilize common traits to build models for lifespan. Predictor traits with model importance of 0.02 or higher used to build Random Forest models of DGRP trait dietary response of (A) mean lifespan, (B) median lifespan, (C) maximum lifespan, (D) initial mortality (α) and (E) rate of aging (β) for mortality from day 21 to the day 95% were dead, (F) variance in day of death, (G) body weight (mg), (H) triglyceride levels (μg) normalized by weight (mg), and (I) glucose levels (μg) normalized by weight (mg).

Many of these traits have previously been associated with the model trait found here. The metabolites kynurenine, quinolinate, myristic acid, and threonine were each used to build at least one lifespan random forest model and have been previously implicated in lifespan.

We visualized the models and their important traits using a combined Kamada-Kawai and forceatlas force-based network organization, which grouped traits in similar clusters based on the push of common factors^13,14^. These networks were further augmented by the addition of top GWAS candidate genes, based on p-value. As expected, the lifespan response traits were grouped and had a few common predictors and genes, specifically late-life alpha and late-life beta, and mean and median lifespans had significant overlap with each other (**Figure 4, Supplementary Figure 4A, and Supplementary Table 3**). For example, myristic acid, glycerol phosphate, and the gene *pbl*, were included in the random forest models for mean and median lifespan or were GWAS candidates, respectively. Similarly, climbing-related response traits clustered, meaning they were used in the same random forest models, though there were more common GWAS candidates, particularly in the AL group (**Supplementary Figure 3A&4B**). Visualizing the connections between these three datasets in this manner gives a better sense of how lifespan and healthspan response traits are built and the genetics that underlie them (**Supplementary Figures 3&4**).

**Figure 4.**
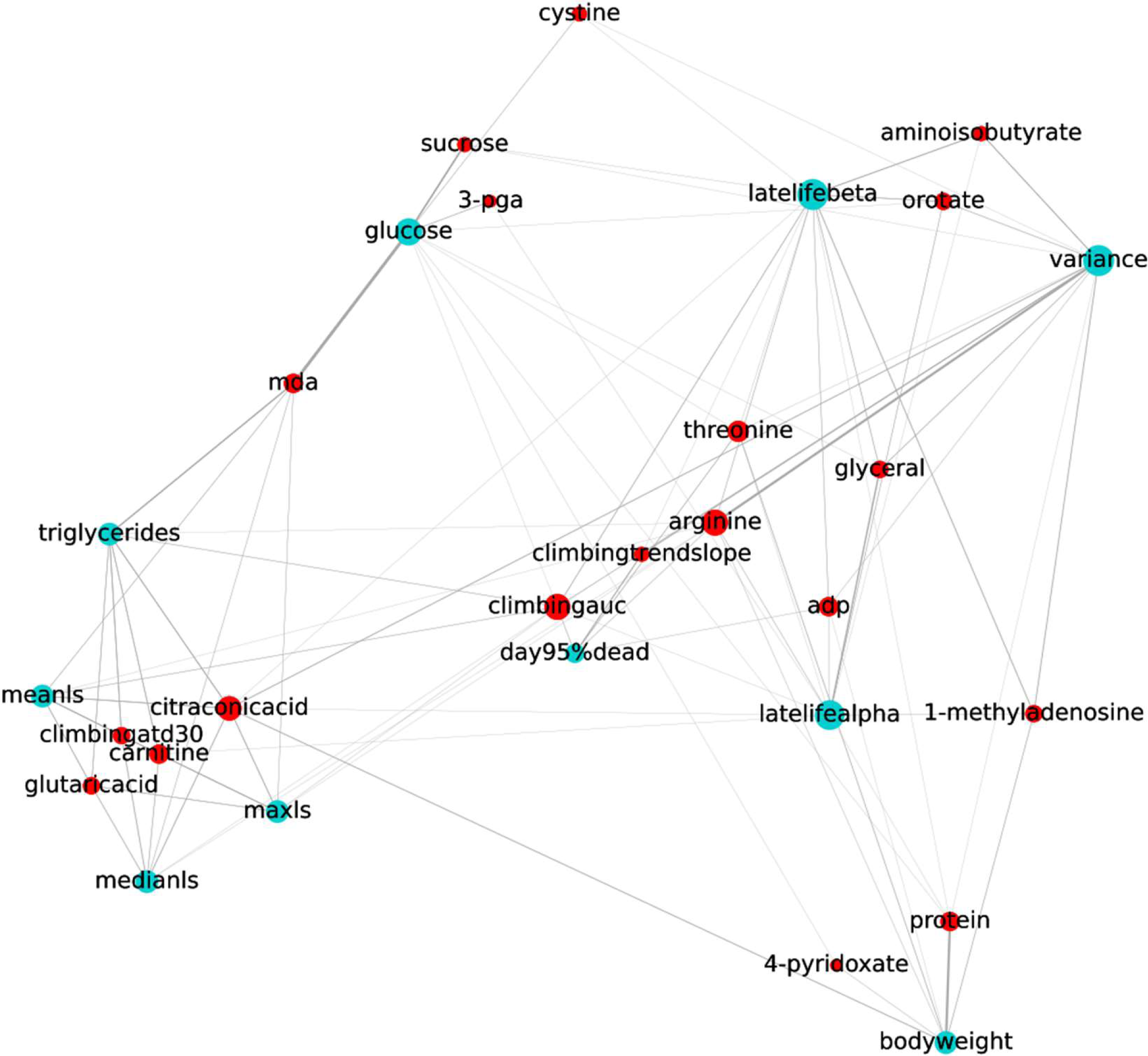
Predictor traits used to build RF lifespan models grouped similar lifespan and healthspan response traits in a network diagram of dietary response. A network diagram of lifespan and healthspan response traits (teal nodes) connected to their top 3 predictor traits with model importance of 0.02 or higher (red nodes) by grey edges whose widths represent their importance. Nodes repel each other, forcing response traits built with similar predictor traits to be pushed together (cluster).

### Mendelian randomization revealed associations between ten metabolites identified through Random Forest and human healthspan and lifespan

We next used the networks developed above to identify multiple response models, with traits identified to be relevant to humans via Mendelian randomization selected for validation in the fly. We explored the utility of the identified metabolite associations from the random forest model approach by applying them in two-sample Mendelian randomization using the IVW method. We identified 10 metabolites in *Drosophila* that were also detected in the large available GWAS for serum metabolites in humans and influence multiple lifespan and healthspan traits (**Figure 5A**). We found that two of the Mendelian randomization-prioritized metabolites, orotate, and choline, were implicated in both healthspan and lifespan random forest models. Both metabolites had clear effects upon longevity, Body Mass Index (BMI), and Frailty Index (P < 0.05/10 = 0.005). Four of the 10 metabolites (urate, inosine, histidine, and threonine) were detected in the lifespan-only random forest models. Urate, inosine, and threonine had significant effects upon longevity (P < 0.05/10 = 0.005) with histidine having significant effects on BMI and Frailty Index (P < 0.05/10 = 0.005) but not longevity (P > 0.05). Amongst the 4 remaining metabolites (quinolinate, glucose, uridine, and kynurenine) detected in the healthspan random forest model, only glucose was significantly associated with longevity using the inverse-variance weighted method (βIVW, 0.005; 95% CI, 0.002–0.008; P = 1.12 × 10^−3^). The remaining metabolites exhibited non-significant associations with longevity but suggestive or significant Frailty Index-related effects. For quinolinate, glucose, and uridine no evidence was found for BMI-related effects (P ≥ 0.1). These genetic associations and their effects can also be characterized by their pleiotropy which was observed in our Mendelian randomization analyses (**Supplementary Table 4**).

**Figure 5.**
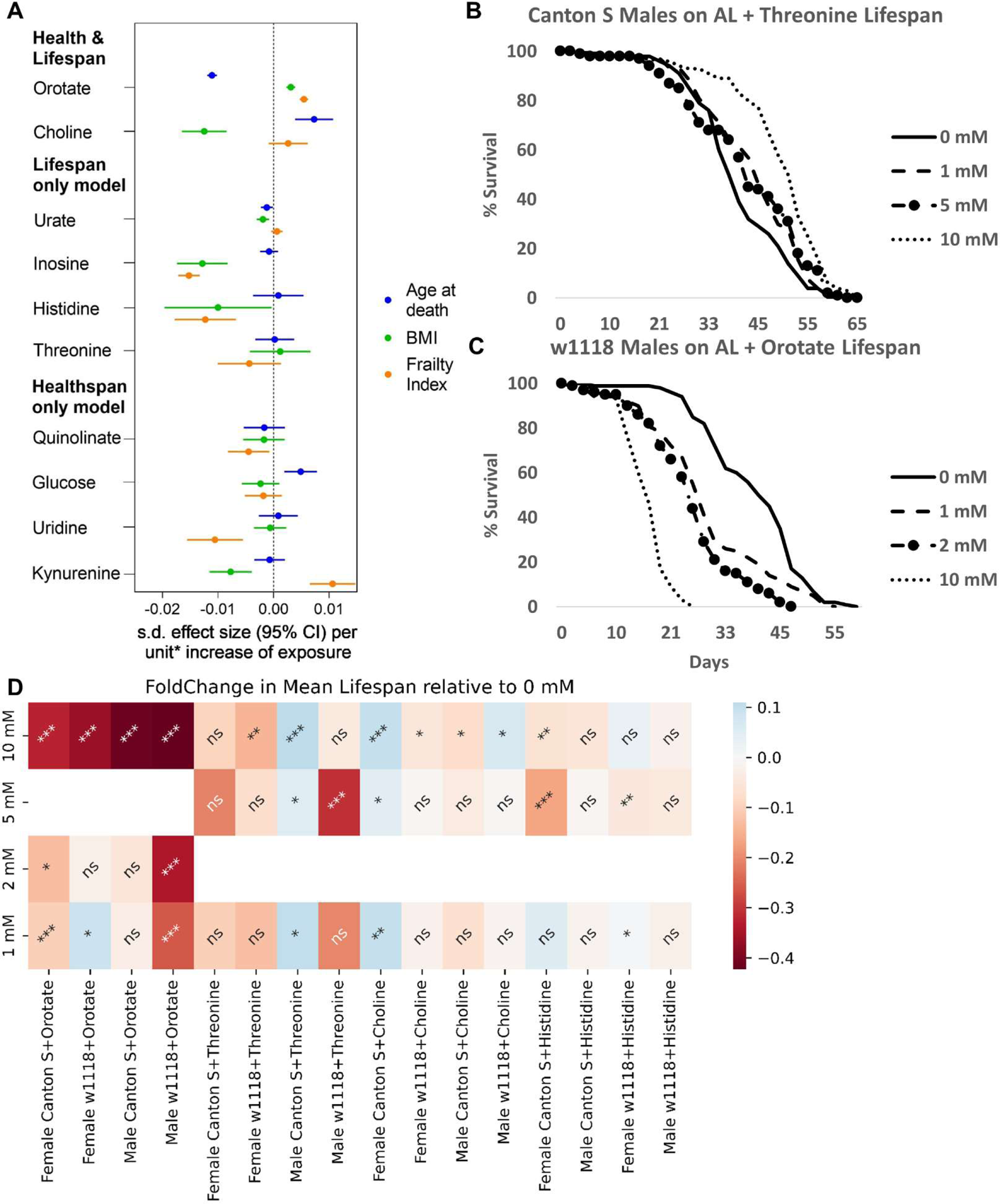
Supplementation of Mendelian randomization and RF overlapping metabolite choline significantly increases fly lifespan in a genotype-, dose-, and sex-specific manner. (A) Mendelian randomization results for metabolites found important in Random Forest models of age at death, BMI, and frailty phenotypes measured in humans. (B) AL diet supplementation of 1-, 5-, or 10-mM threonine significantly extends lifespan in Canton S male Drosophila melanogaster (p ∼ 2.85*10-2, 1.95*10-2, and 9.98*10-10, respectively). (C) AL diet supplementation of 1-, 2-, or 10-mM orotate significantly shortens lifespan in w1118 male Drosophila melanogaster (p ∼ 5.00*10-6, 2.69*10-17, and 8.90*10-30, respectively). (D) Heatmap of mean lifespan fold change values in mean lifespan for males and females of two Drosophila strains on an AL diet supplemented with one of three concentrations of either orotate, threonine, choline, or histidine, compared to the water-only vehicle control (0 mM). Lifespans began with 200 flies per group, with Cox proportional hazards analysis p-values being summarized (ns – not significant, * <= 0.05, ** <= 0.005, *** <= 0.0005). Orotate and choline supplementation repeated twice, threonine and histidine once.

### Shared metabolites between Random Forest lifespan models and Mendelian randomization exhibit sex and strain-specific effects on lifespan in Drosophila melanogaster

To validate the metabolites that were both used to build Random Forest models of lifespan response and found to be relevant for humans via Mendelian randomization, we supplemented our AL diet with 3 concentrations of candidate metabolites. Validated metabolites were selected based on the number of lifespan models they were incorporated in, the average odds-ratio from Mendelian randomization, and the novelty of the lifespan association (**Supplementary Table 5**). The impact of candidate metabolites choline, threonine, histidine, or orotate, on the survival of males and females from two fly strains, w1118 and Canton S, was tracked. Each metabolite showed a strain-, sex-, and dose-specific effect on lifespan (**Figure 5B-D, Supplementary Figure 5**). Orotate supplementation significantly shortened lifespan at 10 mM in both strains and sexes, while choline and threonine extended or shortened lifespan in a strain-, sex-, and dose-specific manner (Fold changes compared to 0 mM controls and Cox proportional hazards p-values summarized in **Figure 5D, Supplementary Figure 5**). Further, 10 mM supplementation of orotate in a DR diet blocked the lifespan effect (**Figure 6A&B**). Supplementation of a DR diet with threonine, choline, or histidine showed similar strain- and sex-specific lifespan effects compared to AL (**Supplementary Figure 6**). Together, these results validate our modeling approach and identify specific metabolites that are relevant for lifespan and have translatable effects in humans.

**Figure 6.**
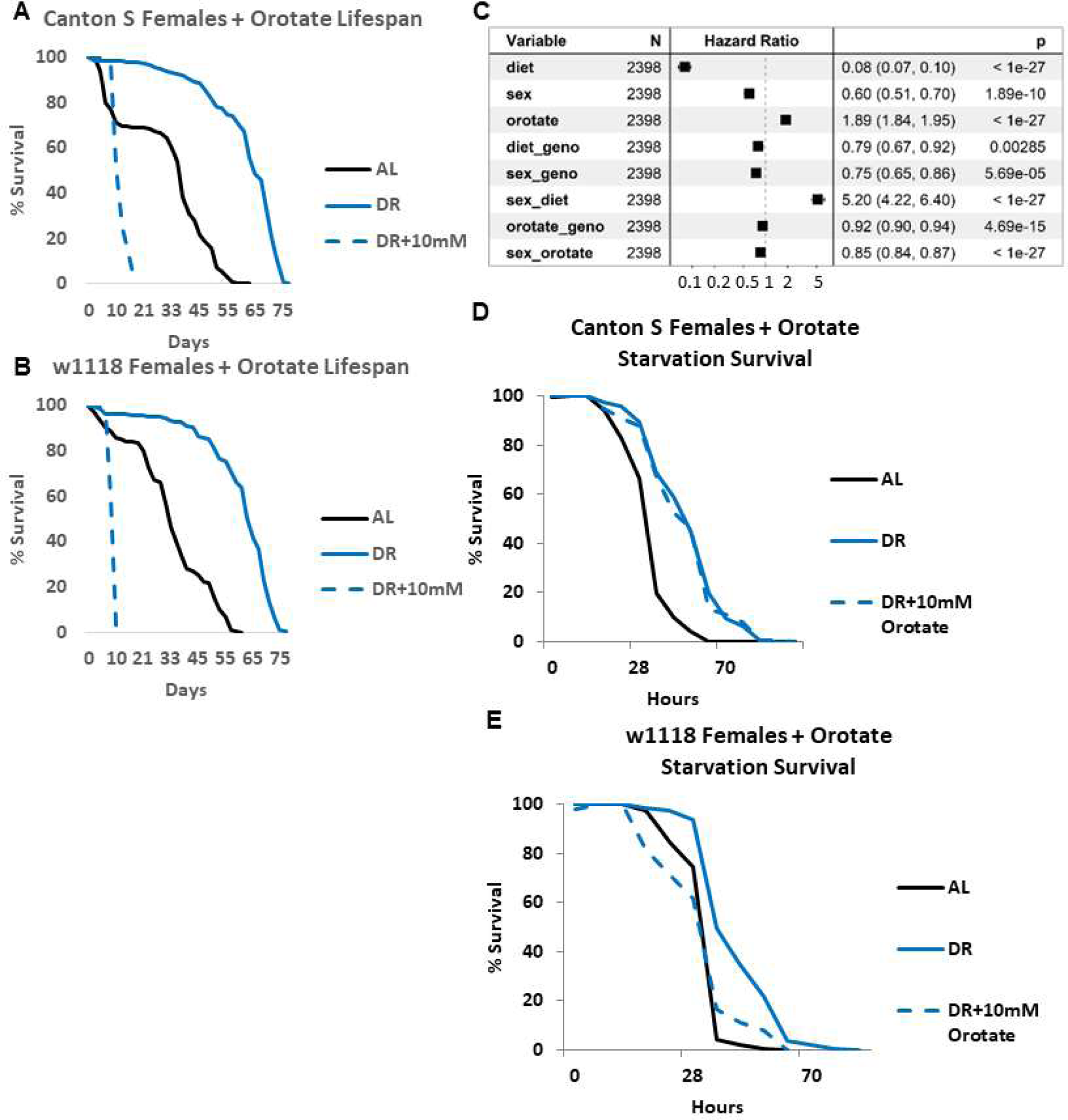
Orotate’s significant decrease in fly lifespan is independent of diet, but the mechanism is strain- and sex-specific. (A) AL diet and DR diet supplemented with 10-mM orotate significantly shortened lifespan compared to DR diet in Canton S female Drosophila melanogaster (p ∼ 3.45*10-52, and 3.02*10-25, respectively). (B) AL diet and DR diet supplemented with 10-mM orotate significantly shortened lifespan compared to DR diet in w1118 female Drosophila melanogaster (p ∼ 1.69*10-50, and 5.66*10-32, respectively). (C) Cox proportional hazards analysis significant results from testing diet (AL as reference), sex (female as reference), orotate levels (0mM as reference), genotype (w1118 as reference), and their interaction terms in flies. (D) AL diet, but not DR diet supplemented with 10-mM orotate, significantly shortened survival under starvation conditions compared to DR diet in Canton S female Drosophila melanogaster (p ∼ 2.83*10-9, and 0.48, respectively). (E) AL diet and DR diet supplemented with 10-mM orotate, significantly shortened survival under starvation conditions compared to DR diet in Canton S female Drosophila melanogaster (p ∼ 2.09*10-4, and 2.00*10-5, respectively), but are not significantly different from each other (p ∼ 0.49). Lifespans began with 200 flies per group, with Cox proportional hazards analysis p-values being summarized (ns – not significant, * <= 0.05, ** <= 0.005, *** <= 0.0005). Orotate supplementation lifespans repeated twice, starvation lifespan once.

### Orotate’s lifespan-shortening effect is diet-independent, but the mechanism is strain- and sex-specific

Using Cox proportional hazards analysis, we dissected the effect sizes for diet (AL was reference), sex (Female was reference), strain (w1118 was reference), and orotate supplementation (0mM was reference) on mortality in the fly (**Figure 6C**). Being on the DR diet independently decreased the hazard ratio (HR=0.08, P < 1 x 10^-27^), being male also independently decreased the hazard ratio (HR=0.60, P = 1.89 x 10^-10^), and 10 mM orotate increased the hazard ratio (HR=1.89, P < 1 x 10^-27^). The fly strain was not significant independently but had significant interaction effects separately with diet, sex, and orotate concentration (HR=0.79, P = 0.0029; HR=0.75, P = 5.69 x 10^-5^, and HR=0.92, P = 4.69 x 10^-15^, respectively). The strongest effect size was the interaction between sex and diet (HR=5.20, P < 1 x 10^-27^). There was also a slight decrease in hazard ratio due to the interaction between sex and orotate concentration (HR=0.85, P < 1 x 10^-27^).

Considering the sex- and strain-specific effects of orotate on lifespan and given orotate’s previously shown role in fatty acid oxidation and lipid metabolism, we used a starvation assay to ascertain whether orotate supplementation was killing the flies by impacting their fat utilization^15,16^. Unlike the survival assays previously mentioned, orotate supplementation of the DR diet prior to starvation conditions did not affect survival compared to the DR control group for Canton S females (**Figure 6D**, ALvsDR P = 2.83 x 10^-9^, ALvsDR+10mM P = 1.79 x 10^-7^, DRvsDR+10mM P = 0.48), but prevented the benefit of the DR diet for w1118 females (**Figure 6E**, ALvsDR P = 2.09 x 10^-4^, ALvsDR+10mM P = 0.49, DRvsDR+10mM P = 2.09 x 10^-5^). The males of each strain showed similar trends, though Canton S males showed no effect of diet on survival under starvation conditions (**Supplementary Figure 6, Supplementary Table 6**). These data show a strain- and sex-specific response to orotate supplementation and suggest separate mechanisms causing the lifespan decreases.

## DISCUSSION

Genetic variability has been shown to greatly influence a strain’s response to various conditions, including changes to diet^4,17,18^. Given that DR is one of the most robust ways to impact lifespan and other healthspan traits across species, understanding the underlying genetic variation and its role in diet-specific responses would help identify the mechanisms by which it functions^19^. While studies have investigated similar connections^4,20–23^, none have employed a comparable systems biology approach that incorporates data from such a diverse range of model organism genotypes. Additionally, our utilization of Mendelian randomization with human data to uncover pertinent outcomes, along with our distinct modeling and visualization strategy for candidate selection and validation, sets our approach apart. Here, we have demonstrated the variability in outcome responses on one of two diets and the differing responses of each genotype. We have demonstrated that trait levels do play a role in different lifespan and healthspan outcomes, despite the lack of correlation across datasets between metabolites and phenotypes. This lack of correlation could indicate non-linear and/or indirect relationships between traits that would not be found in standard correlation analysis. We used a machine learning approach to identify non-linear relationships between our response and predictor traits. Our approach produced a value for each trait based on the proportion of trees/estimators that the trait was included in. This allowed us to confirm the importance of traits previously linked with their response variable, despite the lack of correlation across datasets. By using an unbiased, data-driven approach, we were able to identify relationships that would not be found in standard correlation analysis.

Some examples of traits used to build our response models include the metabolite malondialdehyde, which has previously been linked to glucose and triglyceride levels^24–26^. We found it was important for modeling glucose levels on AL and DR, separately, as well as the DR-AL response in glucose and triglyceride levels. Another metabolite previously associated with triglyceride levels, glyceraldehyde, was also found to be an important feature in our DR-specific model. Additionally, this metabolite was important for our initial mortality and rate of aging dietary response models, aligning with studies showing glyceraldehyde and the enzyme GAPDH as associated with lifespan^27,28^. The metabolite methylhistidine, renowned for its correlation with the extension of lifespan induced by the drug RU486’s transgene activation in flies, as well as its link to mortality and frailty, displayed significance in our initial mortality model under dietary restriction (DR) conditions.^29^ Moreover, it emerged as notably significant in the dietary response influencing maximum lifespan. A downstream metabolite in the well-known kynurenine/NAD+ pathway, quinolinate, which has previously been associated with aging, was an important feature in lifespan models from both diets, separately, and the dietary response^30,31^. Metabolites threonine, arginine, and choline were also included in our lifespan and healthspan models and had previously been reported to influence lifespan^32–40^. Threonine was used to build 8 of the 9 lifespan models and has been shown to increase lifespan in *C. elegans*^38,41–43^. While it has been found that increasing threonine through supplementation or preventing its catabolism can extend lifespan, we found diet-, strain-, and sex-specific effects of threonine. Kynurenine and quinolinate have both been positively associated with age in humans, with quinolinate also being strongly associated with frailty and mortality^44^. Additionally, myristic acid is associated with maximum lifespan potential in mice^45^. The use of traits already shown to play a role in lifespan validates our modeling and suggests the importance of the other candidates it identified. Surprisingly, while climbing and lifespan traits were previously shown to not be correlated, we found climbing traits to be important in our models of lifespan^18^.

We validated our approach to identifying underlying traits implicated in lifespan and healthspan by using traits that have already been shown to play a role in lifespan. While analyses of dietary influences on lifespan can be challenging due to genetic associations characterized by pleiotropy, we used methods such as Mendelian randomization to screen the functional consequences of different genotypes in candidate human aging genes and traits in *Drosophila*. This approach holds the potential to overcome previous shortcomings of both fields and identify potential therapeutic pathways and metabolomic targets for diet response, lifespan, and healthspan. Indeed, we found several traits, including orotic acid, previously unconnected to lifespan or healthspan traits that make promising candidates as potential regulators or biomarkers. Administration of orotic acid is a well-known inducer of fatty liver conditions in rats^44–46^. Although the mechanisms underlying orotic acid-induced fatty liver are not entirely conclusive, various studies have proposed several potential pathways, including impaired fatty acid oxidation, upregulated lipogenesis, and a decrease in hepatic lipid transport^47,48^. In mammalian model organisms, orotic acid-induced fatty liver is species-specific, and thus far, only rats have exhibited susceptibility to its effects^49^. Our results also showed a genotype-specific response to orotic acid supplementation indicative of impaired fatty acid oxidation during starvation in *Drosophila melanogaster*. Nevertheless, the precise molecular mechanism by which orotic acid leads to fat accumulation remains unclear, particularly in the rat liver. It is worth noting that the primary dietary source of orotic acid for the average adult is milk and dairy products (approximately 0.005% of total solids), a concentration insufficient to elicit hepatic changes in rats^50^. However, considering the levels of orotic acid present in numerous dietary and health supplements widely available, as well as recent findings showing increased orotate levels to be detrimental to bone health, there is a need for caution regarding potential health implications for humans^51^.

The few GWAS candidates that are shared between the traits used to build our models could be used to influence multiple health outcomes. The diet-specific nature of our models points to differing mechanisms by which the fly responds to diet. While our data shows the diet-specific impact of metabolites, the mechanisms through which these metabolites work in humans may be different. Studies conducted using summary statistics for Mendelian randomization analysis preclude further stratification by sex or age. Furthermore, studies conducted in populations of predominantly European ancestry also limit the generalizability of findings to other populations.

Considering we saw different responses to metabolite supplementation between both sexes of two *Drosophila* strains, it is expected that there will be similar variation in the response of individual humans to specific metabolites. Ways to optimize the potential effects of traits identified by our approach would be to combine treatments that target specific metabolites based on the genotype of an individual. In total, our work demonstrates the importance and value of incorporating multiple data sets to understand how nature “built” systems that influence lifespan and healthspan traits. Our approach also identifies several new potential mechanisms for how DR influences lifespan and healthspan across multiple genotypes. We demonstrate that combining human and *Drosophila* genetics is an effective approach to further our understanding of the underlying processes regulating longevity and may ultimately contribute to anti-aging strategies in humans.

## Acknowledgements

T.A.U.H. was supported by NIH and NIA award F31AG062112 and is currently supported by NIH/NIA training grant T32AG000266-24. K.A.W. was supported by NIH and NIA award F31AG052299 and NIH/NIA training grant T32AG000266-23. This work was funded by grants from the American Federation of Aging Research (R.B.B. and P.K.); NIH grants R56AG038688, R21AG054121, RO1AG045835 (P.K.), AG049494 (D.P.) and the Larry L. Hillblom Foundation. We thank the Bloomington Drosophila Stock Center, and the Vienna Drosophila Stock Center for providing the flies used.

## Author Contributions

T.A.U.H., V.P.N., K.A.W., R.B.B., and P.K. designed research; T.A.U.H., V.P.N., and E.C. performed research; T.A.U.H. and V.P.N. analyzed data; T.A.U.H., V.P.N., and T.O. provided R and Python code; T.A.U.H., V.P.N., and K.A.W. wrote the paper; R.B.B., J.C., and P.K. provided experimental guidance and funding.

## Materials and Methods

### Fly lines, husbandry, and diet composition

All fly lines were maintained on standard fly yeast extract medium containing 1.55% yeast, 5% sucrose, 0.46% agar, 8.5% corn meal, and 1% acid mix (a 1:1 mix of 10% propionic acid and 83.6% orthophosphoric acid) prepared in distilled water. To prepare the media, cornmeal (85 g), sucrose (50 g), active dry yeast (16 g, “Saf-instant”), and agar (4.6 g) were mixed in a liter of water and brought to a boil under constant stirring. Once cooled down to 60°C 10 ml of acid mix was added to the media. The media were then poured into vials (∼10 ml/vial) or bottles (50 ml/bottle) and allowed to cool down before storing at 4°C for later usage. These vials or bottles were then seeded with some live yeast just before the flies were transferred and used for maintenance of lab stocks, collection of virgins, or setting up crosses.

For each cross, 12-15 virgin females of either *w1118* or *Canton S* strains were mated with 3-5 males of the same genotype in bottles containing an intermediate diet with 1.55% yeast as a protein source. Flies were mated for 5 days, then were removed. 9 days later, non-virgin female progeny were sorted onto an AL (standard diet with 5% yeast) or AL diets containing one of three candidate metabolite concentrations. Concentrations for each metabolite were chosen based on a thorough literature review. 4 to 8 vials of 25 mated female flies per vial were collected for each diet, maintained at 25°C and 65% relative humidity, and were on a 12-hour light/dark cycle.

### Lifespan analysis

Flies developed on standard fly 1.5% yeast extract medium were transferred to the necessary diet within 72 hours after eclosion. For survivorship analysis, vials with 25 mated females were transferred to fresh food every other day and fly survival was scored by counting the number of dead flies. Each lifespan was repeated at least once to generate independent biological replicates^4,18,52–54^. We used Cox proportional hazards analysis implemented in the Python package ‘lifelines’ to analyze the significance of the metabolite concentration on survival outcomes compared to vehicle controls (0 mM). We report the probability that B1=0, from fitting the formula phenotype=B1*variable1. Fold changes in mean lifespan compared to 0 mM were calculated using the following formula, (test concentration – 0 mM)/0 mM.

### Starvation assay

For starvation assays, female flies (On AL media until the specified day) were transferred to vials containing 1% agar and deaths were recorded every 4-6 hours, 3 times a day.

### Genome-wide association mapping

We used DGRP release 2 genotypes, and FlyBase R5 coordinates for gene models. As in Nelson et al., 2016, we used only homozygous positions and a minor allele frequency of R 25% to ensure that the minor allele was represented by many observations at a given polymorphic locus^17,18^. The collected phenotype and genotype data were used as input into an association test via ordinary least-squares regression using the StatsModels module in Python^55^. The linear model was phenotype = β1 x genotype + β2 x diet + β3 x genotype x diet + intercept. Nominal p-values denoted as ‘‘genotype’’ in Figure 1A report the probability that β1 ≠ 0, and those denoted as ‘‘interaction’’ report the probability that β3 ≠ 0.

### Principal Component Analysis (PCA)

All DGRP metabolite and phenotype data for strains were first scaled using the StandardScaler function of the sklearn package. Then missing values were imputed based on the mean of all other strains for that trait. PCA was performed on all strains using the PCA and fit_transform functions in the sklearn decomposition package to observe how well the combined metabolome and phenome can separate samples by diet.

### Pearson Correlation Analysis

Pearson correlations between all metabolites and phenotypes was performed on all strains for each diet combination using the cor() function in the stats package. Heatmaps were created for each correlation matrix with the corrplot function, using the ward.D2 hierarchical clustering method to group predictor traits.

### Random Forest Modeling

DGRP metabolite and phenotype data from each diet combination were split into a response variable set composed of 7 lifespan traits [mean lifespan, median lifespan, max LS, day 95% of flies dead, initial mortality (α), rate of aging (β), and variance in day of death] and 3 healthspan traits [glucose (μg)/weight (mg), triglycerides (μg)/weight (mg), and body weight (mg)], and a predictor set containing the remaining traits. Missing values were imputed based on the mean of all other strains for that trait using the SimpleImputer function in the sklearn.impute package^56^. Predictor and response traits were then split into training (75% of data) and test (25% of data) sets using the train_test_split function in the sklearn.model_selection package^56^. Random Forest models were generated using the RandomForestRegressor function in the sklearn.ensemble package using 10,000 estimators^56^. Mean absolute percentage error and R2 values were determined using the mean_squared_error and r2_score functions in the sklearn.metrics package^56^. Feature importances were extracted from the final model.

### Network Diagrams

Network diagrams were created for all diet combinations using the networkx package in Python to visualize and identify common traits or GWAS candidate genes among the Random Forest models for each diet combination^57^. Network layouts were determined by first applying the spring_layout with a k parameter value of 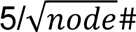, then applying the kamada_kawai_layout with a scale value of 5.

### Mendelian randomization

Genetic instruments (i.e., SNPs) strongly (P<5×10–6) predicting metabolites of interest were extracted from an existing GWAS of serum metabolites in 1960 individuals (96.6% female) of European descent^58^. As the primary outcome of the present analysis, we obtained genetic association estimates for the variants selected as metabolite status instruments with combined parental age at death, from the UK Biobank participants of European descent. Parental lifespan is widely used as an outcome in genetic association studies as offspring inherit one-half of their genetic code from their parents resulting in genotypic and phenotypic correlations^59^.

Secondary outcomes related to health and lifespan included Body Mass Index (BMI) and Frailty Index. Briefly, summary statistics for Frailty Index were collected from a recent GWAS carried out among participants in the UK Biobank^60^. The Frailty Index was constructed based on 49 items ranging from physical to mental well-being and calculated as a proportion of the sum of all deficits. Genetic predictors of BMI were obtained from the largest sex-specific meta-analysis of GWAS in the UK Biobank GWAS and the GIANT consortium^61^. More details of the GWAS on each of the outcomes can be found in their respective published articles.

As the main analysis for Mendelian randomization, we used the inverse-variance weighted method (IVW)^62^ (Supplementary Table 4). IVW is considered to provide the most accurate estimate but is known to be sensitive to pleiotropy^62^. As sensitivity analyses, we also employed the weighted median approach (WM)^63^ and Mendelian randomization-Egger regression^64^ (Supplementary Table 4). The weighted median method assumes >50% of the weight comes from valid SNPs and can generate consistent causal estimates^63^. The Mendelian randomization-Egger regression detects possible pleiotropic effects and provides corrected estimates for pleiotropy^64^. The p-value for the intercept in Mendelian randomization-Egger was used to detect the directional pleiotropic effect^64^. To gauge evidence of directional pleiotropy, we used Cochrane’s Q test^65^, with this being the measure of heterogeneity between ratio estimates of variants (Supplementary Table 4). Finally, to identify candidate genes for functional screening in *Drosophila* we set the significance threshold using Bonferroni correction, at P < 5.00 × 10–3 (= 0.05/10) to declare a causal relationship for the IVW-based Mendelian randomization estimate. Associations with P < 0.05, but not reaching the Bonferroni-corrected threshold, were reported as suggestive of association. All analyses were performed using the 2-sample Mendelian randomization analyses conducted in R using the TwoSampleMR^66^ package.

### Data and Software Availability

https://data.mendeley.com/datasets/67f2sxfwxn/draft?a=6d0eaf2c-385b-498d-a599-7a40211a850e

## Supplementary Figure Titles and Legends

**Supplementary Figure 1. Variation of DGRP dietary response in lifespan and healthspan response traits.** Dietary response (DR-AL) of 160 DGRP strains in 7 lifespan and healthspan metrics, strains plotted in ascending order.

**Supplementary Figure 2. DGRP metabolite and phenotype trait correlations on DR and AL, together (A), or separate (B&C).** Heatmap of metabolite and phenotype value correlations on DR and AL, together, or separate. Traits are clustered via hierarchical clustering using the Ward method, with 5 clusters highlighted on the diagonal. Pearson correlation values are shown as a gradient from 1 (dark blue) to -1 (dark red).

**Supplementary Figure 3. Network diagram of DGRP random forest models of response traits and the predictor traits used to build them; separated by diet group.** Network diagram of response traits (teal nodes) connected to predictor traits with model importance of 0.02 or higher (red nodes) by grey edges whose widths represent their importance. Network layout is a combination of a spring layout followed by a kamada kawai layout, with nodes repelling each other, forcing response traits built with similar predictor traits to be pushed together (cluster). (A) AL, (B) DR, and (C) DR & AL.

**Supplementary Figure 4. Network diagram of DGRP random forest models of response traits, the predictor traits used to build them, and GWAS candidate genes; separated by diet group.** Network diagram of response traits (teal nodes) connected to predictor traits with model importance of 0.02 or higher (red nodes) by grey edges whose widths represent their importance. GWAS candidates (green nodes) with p-values of magnitude 10^-5^ or lower are connected to their trait by edges with widths inversely weighted by p-value (thicker edges represent lower p-values). Network layout is a combination of a spring layout followed by a kamada kawai layout, with nodes repelling each other, forcing response traits built with similar predictor traits to be pushed together (cluster). (A) AL, (B) DR, and (C) DR & AL.

**Supplementary Figure 5. Lifespans of Canton S or w1118 females and males on AL food supplemented with one concentration of a metabolite.** Survival curves for females and males of *Drosophila melanogaster* strains Canton S or w1118 on an AL diet supplemented with one of three concentrations of either choline (A-C), histidine (D-G), or threonine (H-K), or the water vehicle control. All experiments began with 200 flies per group, with cox proportional hazards analysis used to analyze lifespan differences compared to the vehicle only (0 mM) control. Fold changes in mean lifespan compared to the 0 mM control and summary p-values are shown in **Figure 5D**. Orotate and choline supplementation lifespans were repeated twice, and other metabolites once.

**Supplementary Figure 6. Lifespans of Canton S or w1118 females and males on DR food supplemented with 10mM of orotate, threonine, choline, or histidine.** A subset of survival curves for females or males of *Drosophila melanogaster* strains Canton S or w1118 on a DR diet supplemented with 10 mM of either orotate (A-B), threonine (C-D), choline (E), or histidine (F). (G) Heatmap of mean lifespan fold change values in mean lifespan for males and females of two *Drosophila* strains on a DR diet supplemented with 10 mM of either orotate, threonine, choline, or histidine, compared to the water-only vehicle control (DR). Lifespans began with 200 flies per group, with *Cox proportional hazards* analysis p-values being summarized (ns – not significant, * <= 0.05, ** <= 0.005, *** <= 0.0005, **** <= 0.00005). Orotate supplementation lifespans repeated twice, and other metabolites once.

## Supplementary Table Titles and Legends

**Supplementary Table 1. Trait list and DGRP strains with missing values.** List of metabolite and phenotype traits and their descriptions on the first sheet. The second sheet contains a list of DGRP strains with missing phenotype values that were imputed with the mean of all other strains for that trait.

**Supplementary Table 2. DGRP datasets.** Raw DGRP data on each diet used for correlation and modeling analysis.

**Supplementary Table 3. Random Forest response model traits.** Order of traits used to build each random forest response model by the proportion of trees/estimators that trait was included in out of the 10,000 trees/estimators run for each model.

**Supplementary Table 4. Mendelian randomization Results.**

**Supplementary Table 5. Model trait selection and validation determination.**

**Supplementary Table 6. Lifespan Summary Stats and Statistics.** Cox proportional hazards analysis statistics using survival package in Python. Sheets contain statistics for specified experiments. Deaths for each group are at the top of each sheet, with comparisons in the rows below. The last two columns of each sheet contain the comparison statistics and means for that row’s comparison.

