## Supplementary Figures for "Systems biology and machine learning approaches identify metabolites that influence dietary lifespan and healthspan responses across flies and humans"

### Slide 1
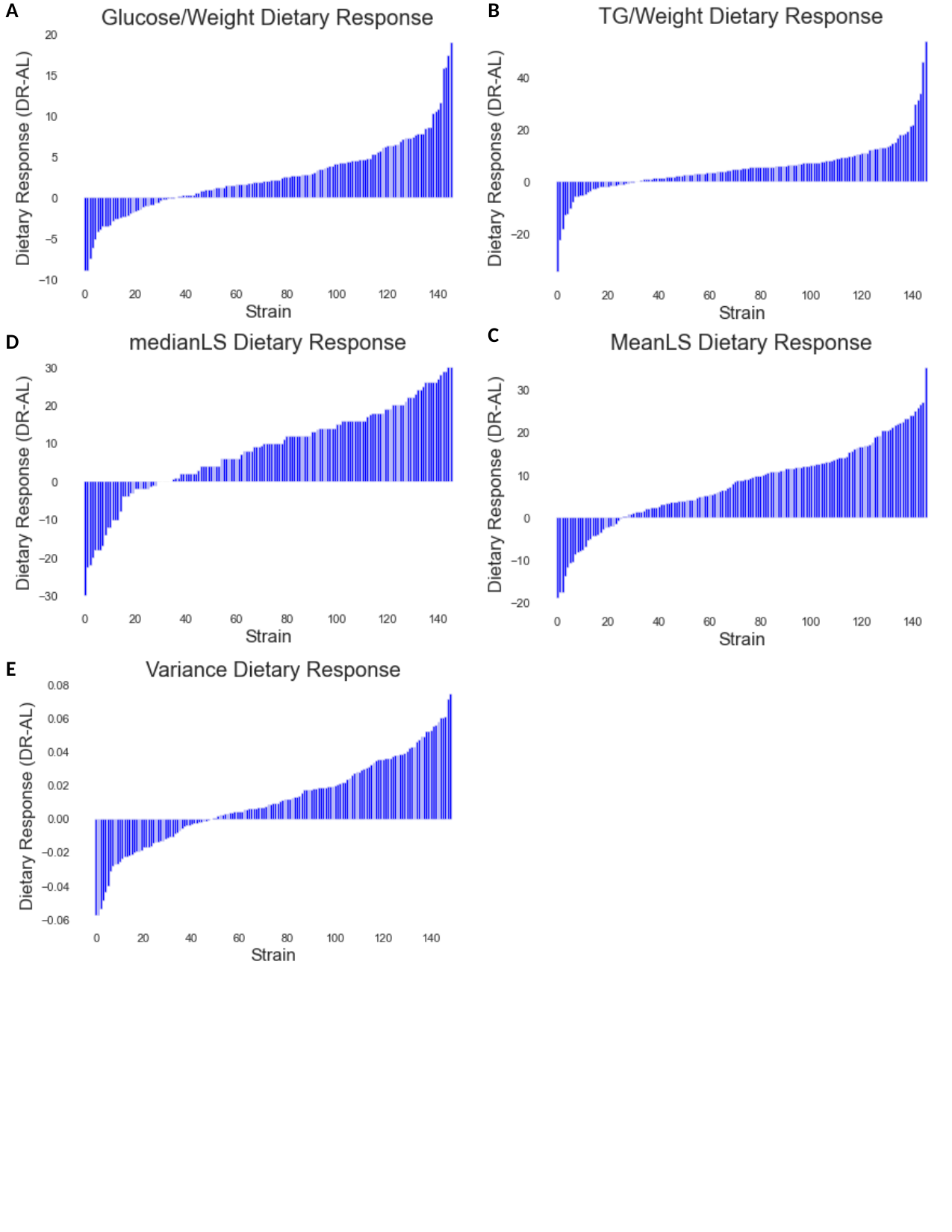

A
B
C
D
E

### Slide 2
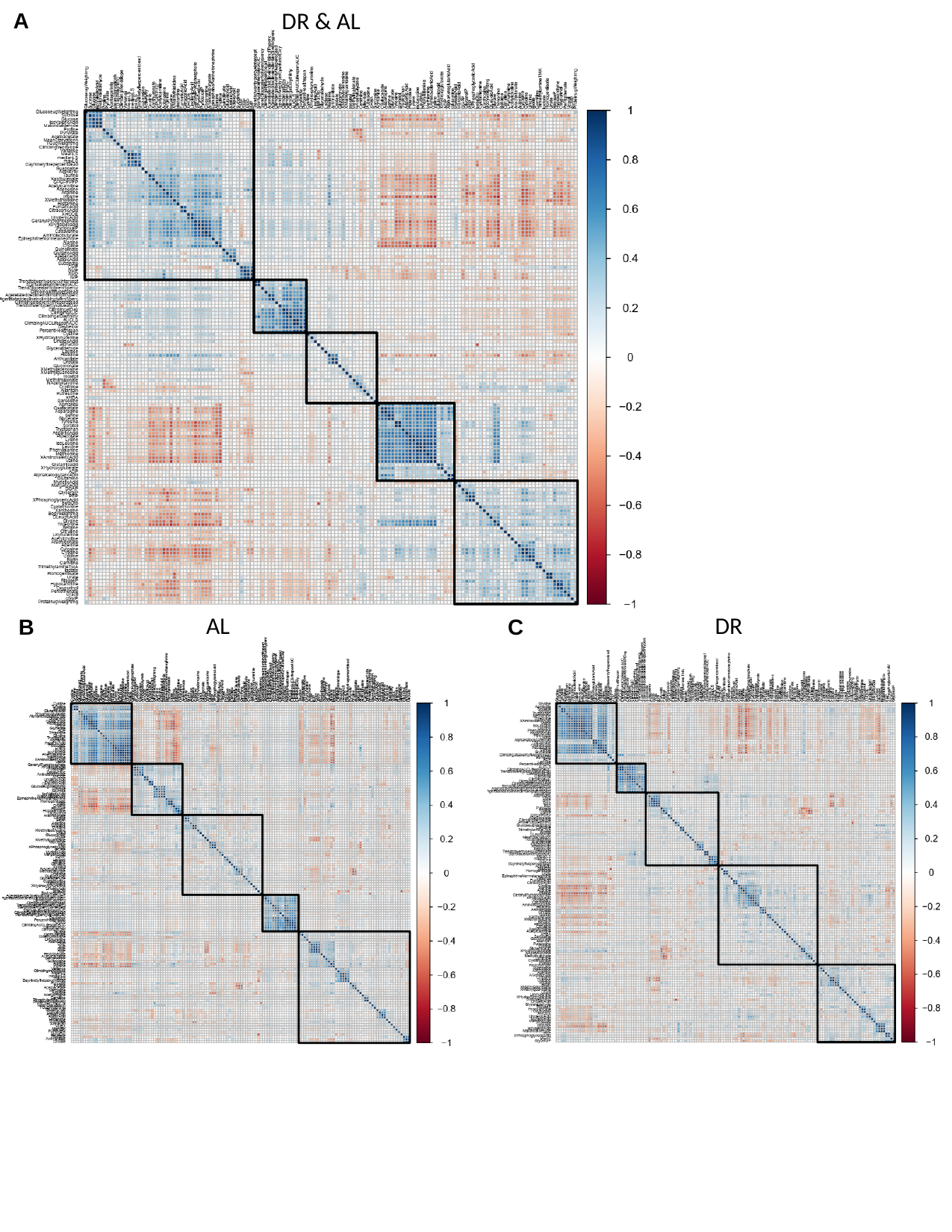

DR & AL
A
AL
DR
C
B

### Slide 3
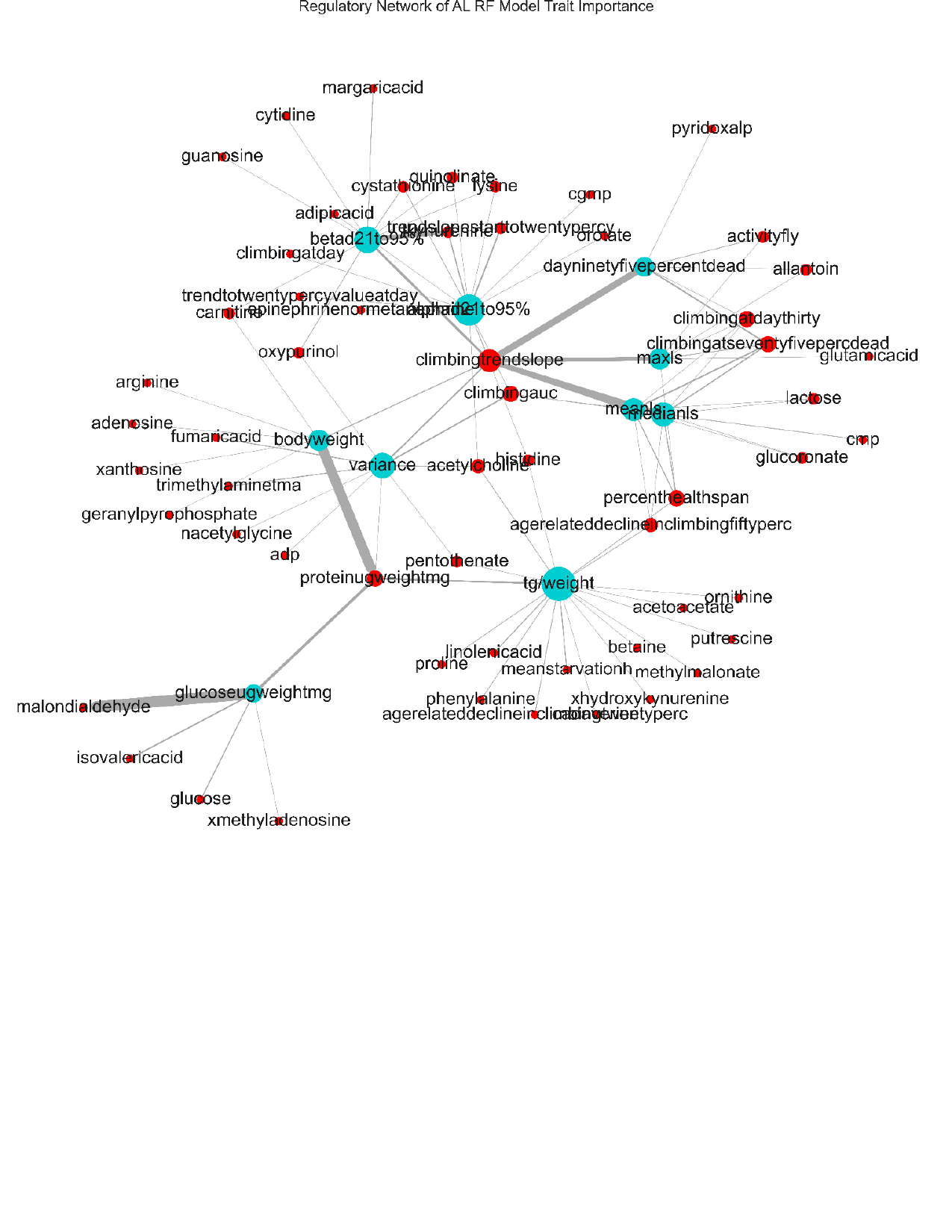

### Slide 4
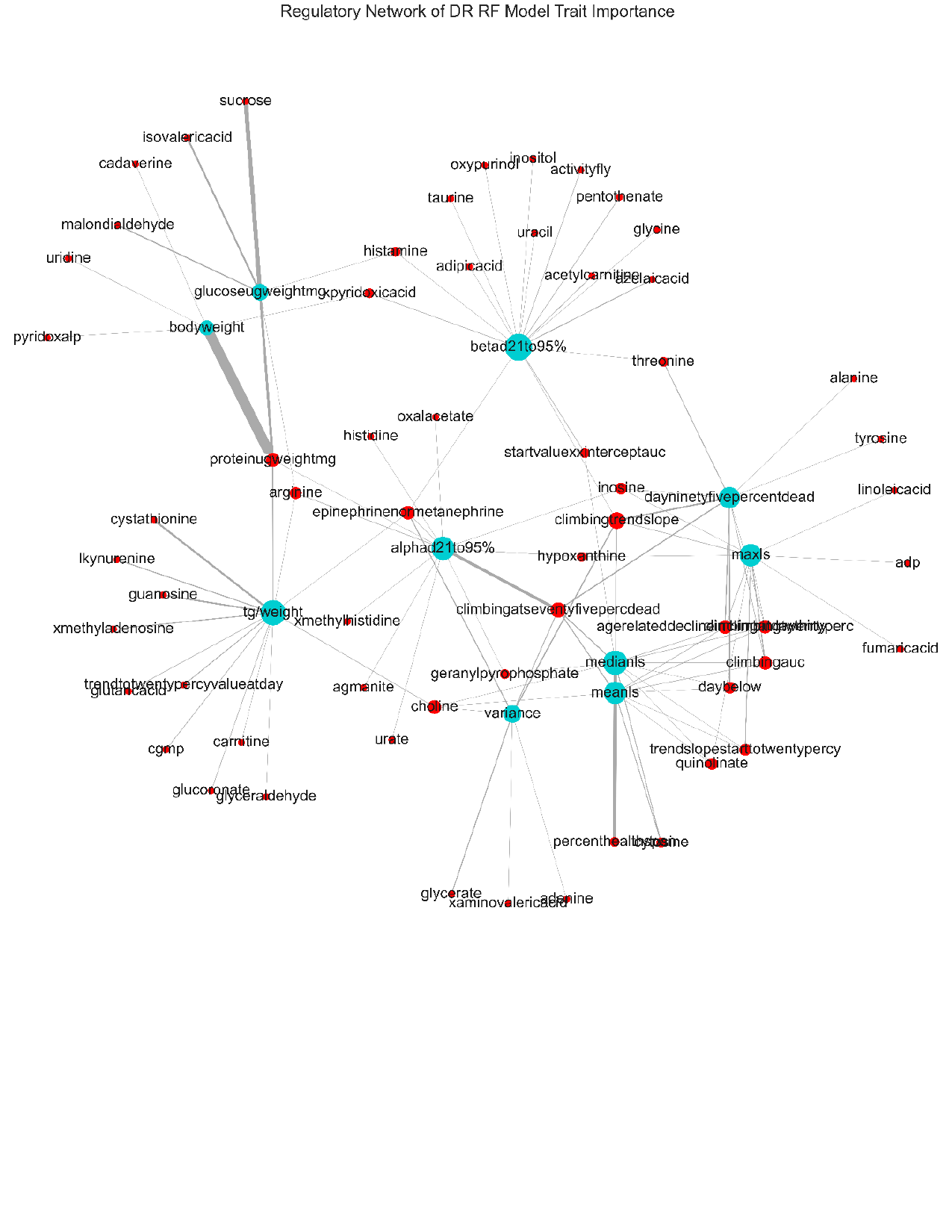

### Slide 5
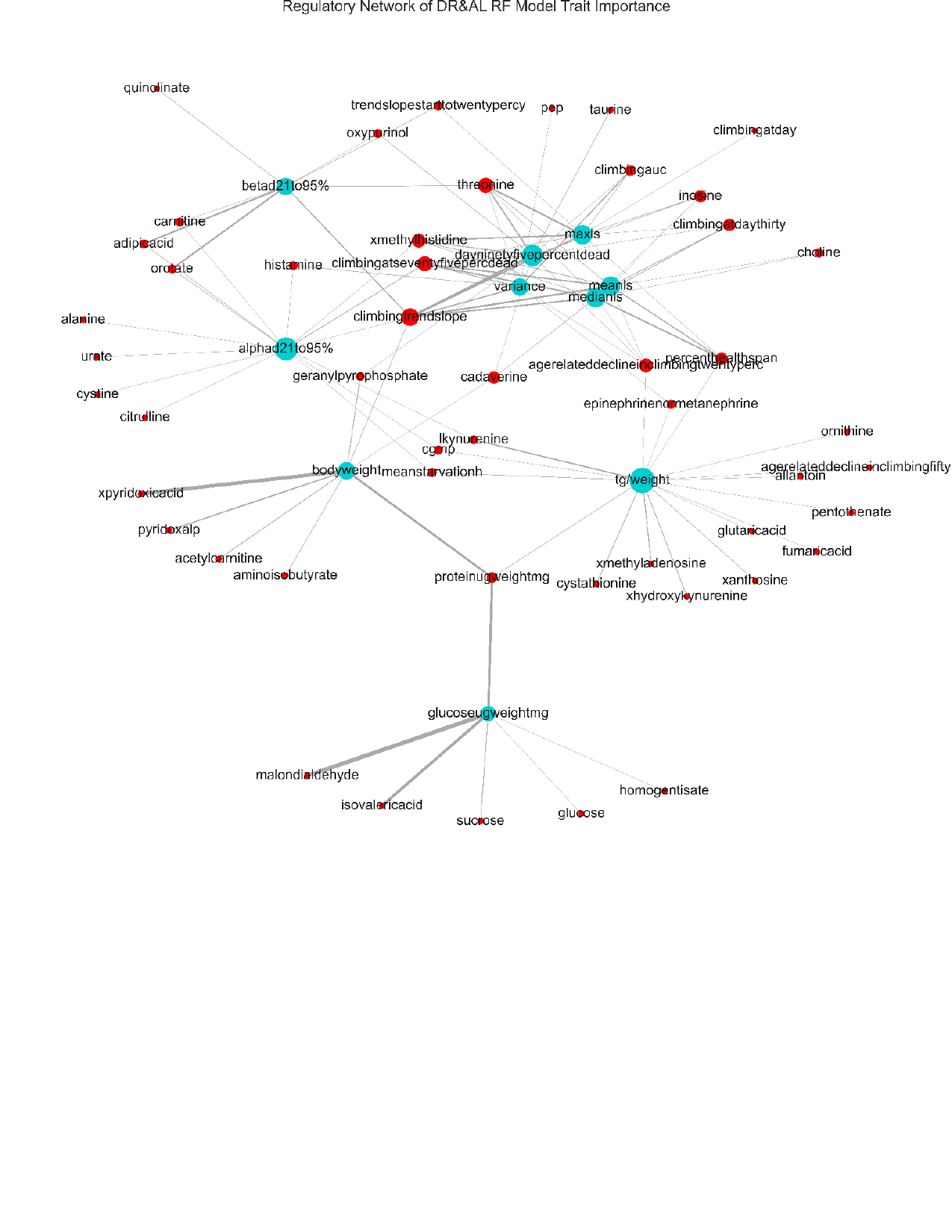

### Slide 6
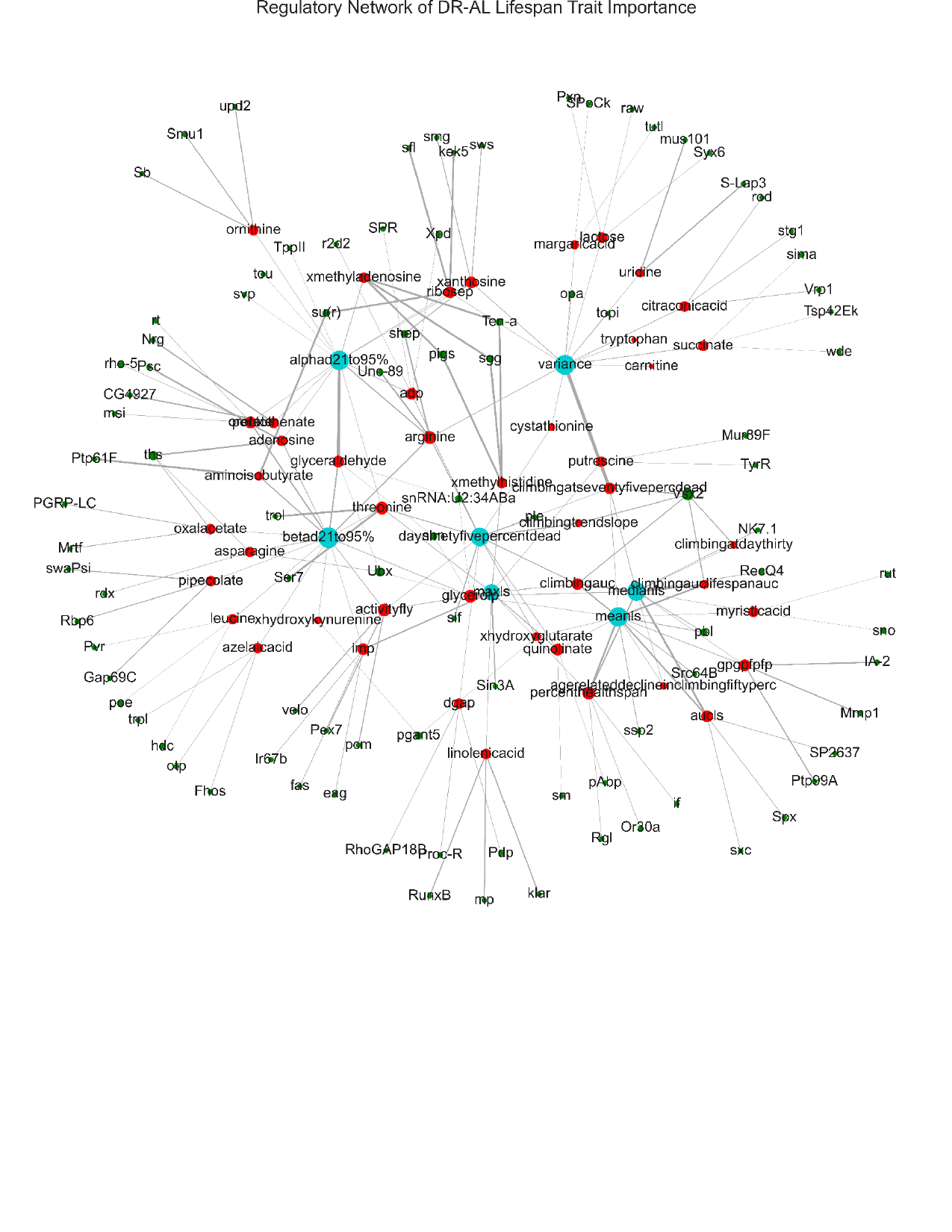

### Slide 7
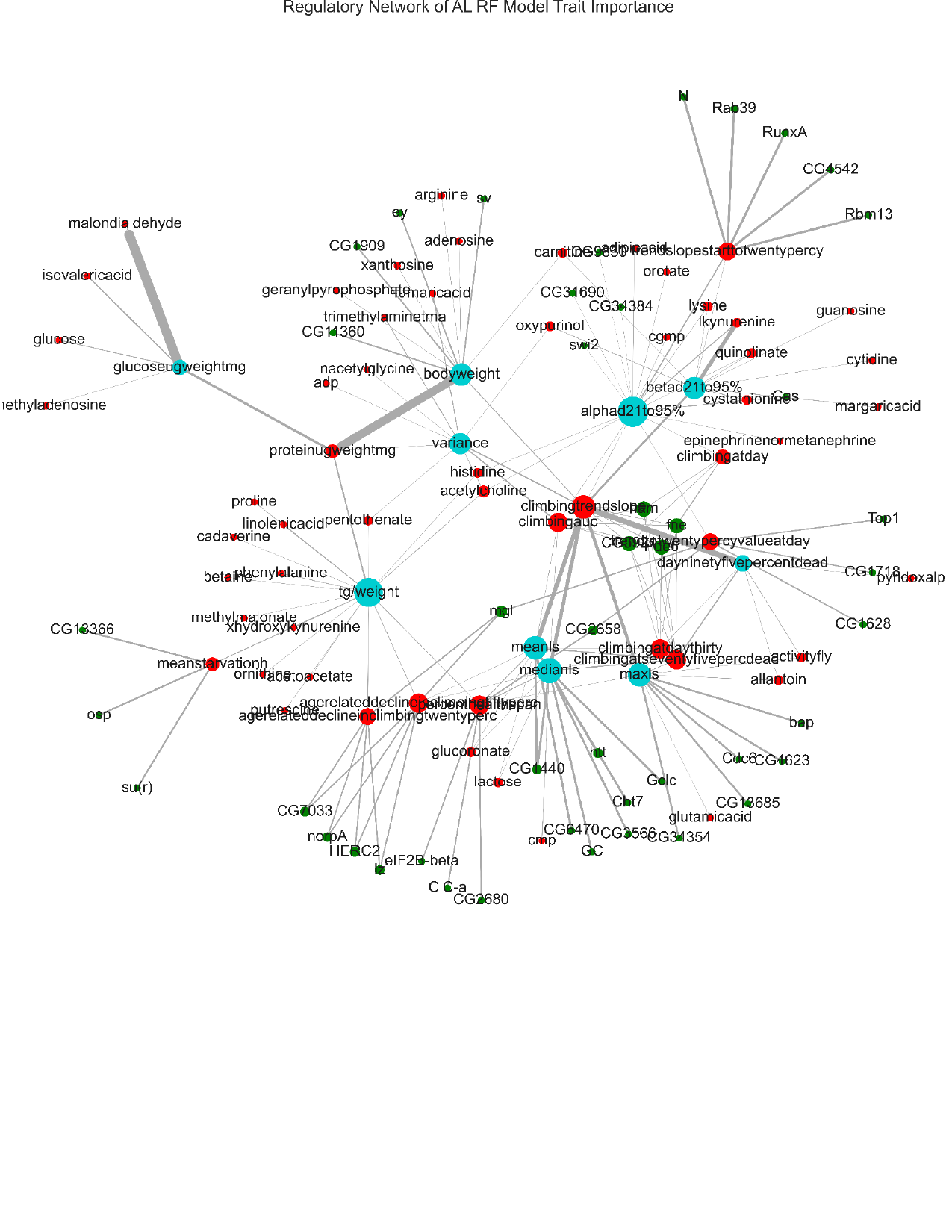

### Slide 8
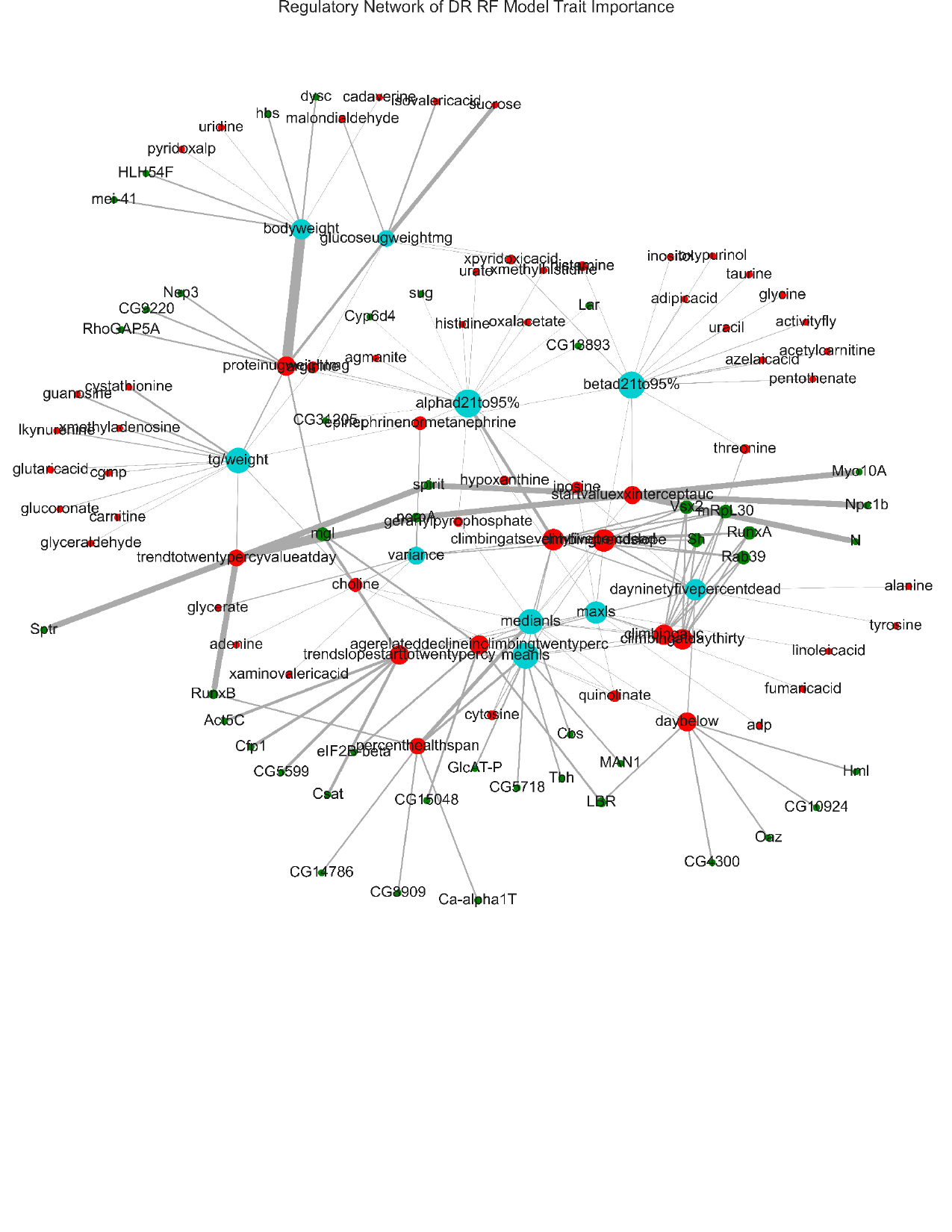

### Slide 9
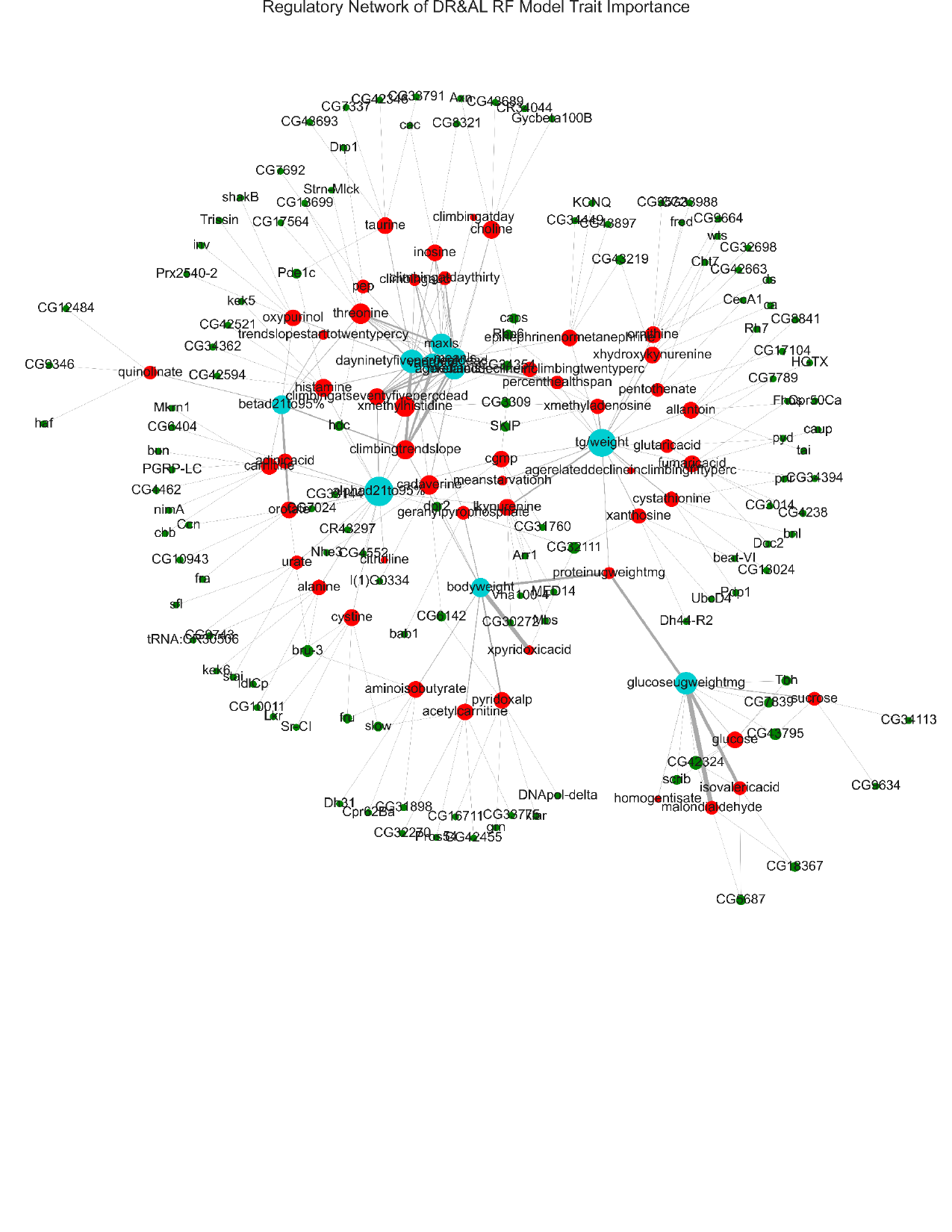

### Slide 10
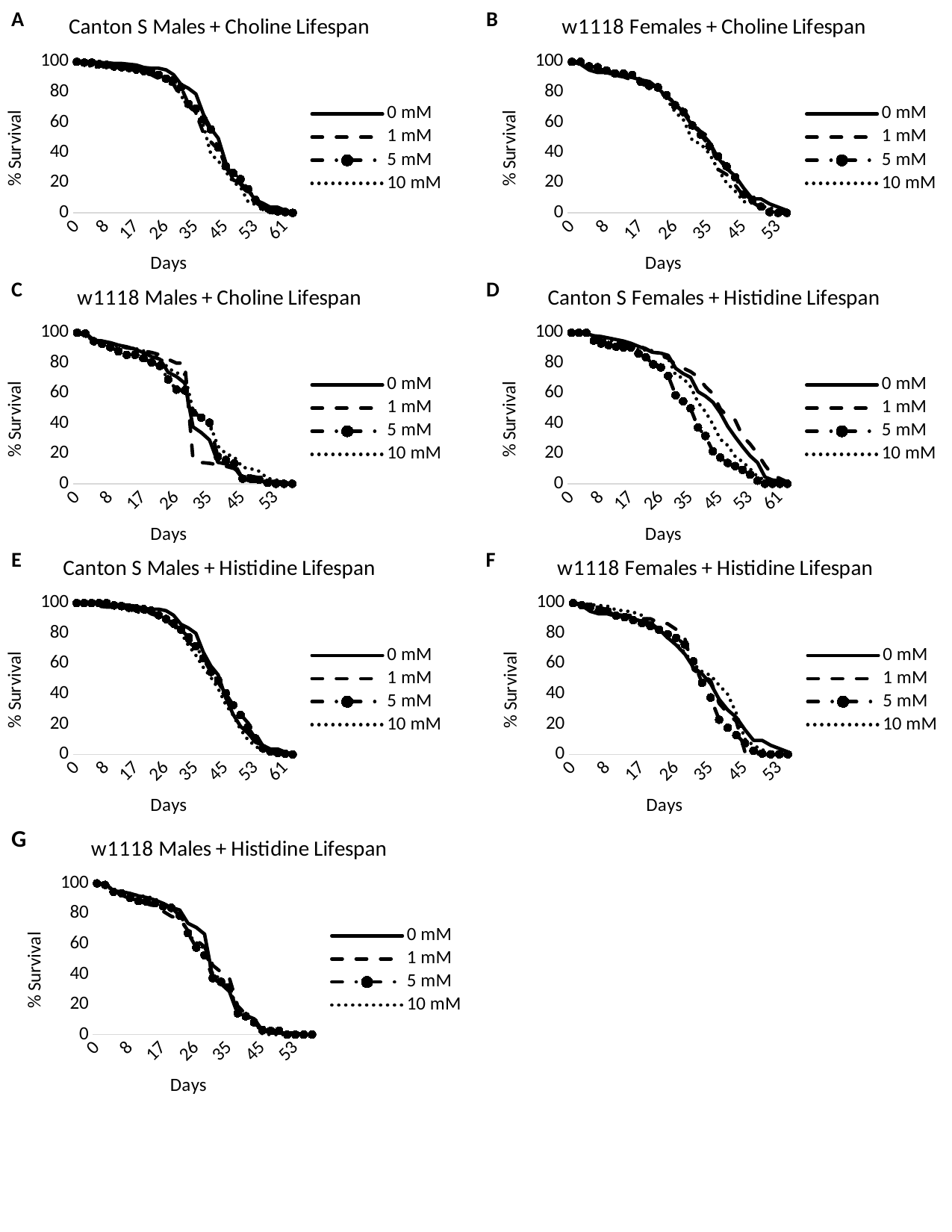

#### Chart: w1118 Females + Choline Lifespan
| Category | 0 mM | 1 mM | 5 mM | 10 mM |
|---|---|---|---|---|
| 0 | 100.0 | 100.0 | 100.0 | 100.0 |
| 2 | 98.3695652173913 | 100.0 | 100.0 | 98.51485148514851 |
| 4 | 94.56521739130434 | 97.51243781094527 | 96.92307692307692 | 96.53465346534654 |
| 6 | 92.93478260869566 | 96.01990049751244 | 96.41025641025641 | 95.54455445544555 |
| 8 | 92.93478260869566 | 93.53233830845771 | 94.35897435897436 | 94.55445544554455 |
| 10 | 91.30434782608695 | 91.04477611940298 | 92.3076923076923 | 92.07920792079207 |
| 12 | 91.30434782608695 | 90.04975124378109 | 92.3076923076923 | 91.58415841584159 |
| 14 | 89.67391304347827 | 88.05970149253731 | 91.28205128205128 | 91.08910891089108 |
| 17 | 88.04347826086956 | 85.57213930348259 | 86.66666666666667 | 88.61386138613861 |
| 19 | 86.95652173913044 | 83.58208955223881 | 84.1025641025641 | 86.63366336633663 |
| 21 | 82.6086956521739 | 80.09950248756219 | 83.07692307692308 | 83.16831683168317 |
| 24 | 77.17391304347827 | 77.11442786069651 | 77.94871794871796 | 75.74257425742574 |
| 26 | 72.28260869565217 | 73.13432835820896 | 71.28205128205128 | 66.83168316831683 |
| 28 | 66.30434782608695 | 68.1592039800995 | 66.66666666666667 | 61.386138613861384 |
| 31 | 57.608695652173914 | 59.20398009950249 | 57.94871794871795 | 48.01980198019802 |
| 33 | 53.26086956521739 | 53.73134328358209 | 51.794871794871796 | 45.54455445544554 |
| 35 | 46.7391304347826 | 50.24875621890547 | 44.1025641025641 | 39.10891089108911 |
| 38 | 36.41304347826087 | 28.855721393034827 | 37.43589743589744 | 28.71287128712872 |
| 40 | 29.347826086956516 | 25.373134328358205 | 30.769230769230774 | 19.80198019801979 |
| 42 | 24.456521739130437 | 18.407960199004975 | 23.58974358974359 | 14.356435643564353 |
| 45 | 16.304347826086953 | 10.945273631840791 | 12.307692307692307 | 7.425742574257427 |
| 47 | 9.23913043478261 | 6.96517412935323 | 8.717948717948715 | 6.435643564356425 |
| 49 | 9.23913043478261 | 3.4825870646766077 | 4.102564102564102 | 4.455445544554465 |
| 51 | 5.978260869565219 | 0.4975124378109399 | 0.512820512820511 | 3.465346534653463 |
| 53 | 3.8043478260869534 | 0.4975124378109399 | 0.0 | 1.9801980198019749 |
| 55 | 1.6304347826086882 | 0.0 | 0.0 | 0.0 |
#### Chart: Canton S Males + Choline Lifespan
| Category | 0 mM | 1 mM | 5 mM | 10 mM |
|---|---|---|---|---|
| 0 | 100.0 | 100.0 | 100.0 | 100.0 |
| 2 | 100.0 | 98.5 | 99.49238578680203 | 99.5 |
| 4 | 100.0 | 98.5 | 99.49238578680203 | 96.5 |
| 6 | 99.46808510638297 | 97.5 | 98.4771573604061 | 96.5 |
| 8 | 99.46808510638297 | 97.5 | 97.96954314720813 | 96.5 |
| 10 | 98.93617021276596 | 97.0 | 96.95431472081218 | 96.5 |
| 12 | 98.93617021276596 | 97.0 | 96.44670050761421 | 95.5 |
| 14 | 98.40425531914893 | 96.0 | 95.93908629441624 | 95.5 |
| 17 | 97.87234042553192 | 94.0 | 94.9238578680203 | 95.5 |
| 19 | 96.27659574468085 | 92.5 | 93.90862944162437 | 94.0 |
| 21 | 95.74468085106383 | 91.5 | 93.40101522842639 | 93.5 |
| 24 | 95.74468085106383 | 89.0 | 91.37055837563452 | 90.0 |
| 26 | 94.68085106382979 | 86.0 | 88.83248730964468 | 88.0 |
| 28 | 91.48936170212767 | 85.0 | 87.81725888324873 | 86.0 |
| 31 | 85.1063829787234 | 76.5 | 83.248730964467 | 77.5 |
| 33 | 82.4468085106383 | 71.0 | 72.08121827411168 | 71.0 |
| 35 | 78.72340425531915 | 66.0 | 69.03553299492386 | 66.0 |
| 38 | 65.42553191489361 | 53.0 | 61.92893401015229 | 58.0 |
| 40 | 55.851063829787236 | 47.0 | 55.32994923857868 | 39.0 |
| 42 | 49.46808510638297 | 42.00000000000001 | 43.147208121827404 | 34.5 |
| 45 | 33.51063829787235 | 30.0 | 30.964467005076145 | 27.5 |
| 47 | 21.808510638297875 | 24.0 | 26.39593908629442 | 20.5 |
| 49 | 19.68085106382979 | 19.0 | 22.33502538071066 | 17.0 |
| 51 | 14.893617021276597 | 12.5 | 15.736040609137063 | 7.5 |
| 53 | 8.510638297872347 | 7.0 | 8.629441624365484 | 5.5 |
| 55 | 6.38297872340425 | 3.5 | 4.060913705583758 | 3.0 |
| 57 | 3.723404255319153 | 1.0 | 2.030456852791872 | 0.0 |
| 59 | 3.723404255319153 | 0.0 | 1.015228426395936 | 0.0 |
| 61 | 1.5957446808510696 | 0.0 | 0.507614213197968 | 0.0 |
| 63 | 0.0 | 0.0 | 0.0 | 0.0 |A
B
#### Chart: w1118 Males + Choline Lifespan
| Category | 0 mM | 1 mM | 5 mM | 10 mM |
|---|---|---|---|---|
| 0 | 100.0 | 100.0 | 100.0 | 100.0 |
| 2 | 100.0 | 97.98994974874373 | 99.43820224719101 | 97.91666666666667 |
| 4 | 95.08196721311475 | 95.97989949748744 | 94.38202247191012 | 94.79166666666667 |
| 6 | 94.53551912568307 | 94.47236180904522 | 92.69662921348315 | 92.70833333333333 |
| 8 | 93.44262295081967 | 92.96482412060301 | 90.4494382022472 | 91.14583333333333 |
| 10 | 91.80327868852459 | 91.4572864321608 | 87.64044943820225 | 91.14583333333333 |
| 12 | 90.7103825136612 | 89.94974874371859 | 85.3932584269663 | 90.10416666666667 |
| 14 | 89.07103825136612 | 89.44723618090453 | 85.3932584269663 | 89.58333333333333 |
| 17 | 86.88524590163934 | 87.93969849246231 | 83.14606741573033 | 88.02083333333333 |
| 19 | 84.15300546448087 | 86.93467336683418 | 80.33707865168539 | 86.45833333333333 |
| 21 | 82.51366120218579 | 85.42713567839196 | 78.08988764044943 | 83.33333333333334 |
| 24 | 73.77049180327869 | 82.41206030150754 | 69.10112359550561 | 76.5625 |
| 26 | 71.03825136612022 | 79.89949748743719 | 62.359550561797754 | 73.4375 |
| 28 | 66.66666666666667 | 79.89949748743719 | 61.79775280898877 | 72.39583333333333 |
| 31 | 37.704918032786885 | 16.08040201005025 | 47.19101123595506 | 48.4375 |
| 33 | 33.87978142076503 | 14.070351758793976 | 43.82022471910112 | 44.270833333333336 |
| 35 | 28.96174863387978 | 13.5678391959799 | 40.44943820224719 | 40.104166666666664 |
| 38 | 14.207650273224047 | 12.562814070351763 | 17.97752808988764 | 25.0 |
| 40 | 13.114754098360663 | 11.557788944723612 | 15.730337078651687 | 19.791666666666657 |
| 42 | 10.928961748633881 | 10.050251256281399 | 13.483146067415731 | 17.708333333333343 |
| 45 | 3.278688524590166 | 5.527638190954775 | 3.3707865168539257 | 10.9375 |
| 47 | 1.639344262295083 | 5.0251256281406995 | 3.3707865168539257 | 9.895833333333343 |
| 49 | 1.639344262295083 | 4.0201005025125625 | 2.8089887640449405 | 8.333333333333343 |
| 51 | 1.639344262295083 | 1.0050251256281513 | 0.5617977528089853 | 3.645833333333343 |
| 53 | 1.639344262295083 | 1.0050251256281513 | 0.0 | 2.604166666666657 |
| 55 | 1.0928961748633839 | 1.0050251256281513 | 0.0 | 1.5625 |
| 57 | 0.0 | 0.0 | 0.0 | 0.0 |
#### Chart: Canton S Females + Histidine Lifespan
| Category | 0 mM | 1 mM | 5 mM | 10 mM |
|---|---|---|---|---|
| 0 | 100.0 | 100.0 | 100.0 | 100.0 |
| 2 | 99.5 | 99.41520467836257 | 100.0 | 100.0 |
| 4 | 99.5 | 99.41520467836257 | 100.0 | 100.0 |
| 6 | 98.0 | 97.07602339181287 | 94.89795918367346 | 97.40932642487047 |
| 8 | 97.5 | 95.32163742690058 | 92.85714285714286 | 96.37305699481865 |
| 10 | 96.5 | 93.5672514619883 | 91.83673469387755 | 95.85492227979275 |
| 12 | 95.5 | 93.5672514619883 | 90.81632653061224 | 93.78238341968913 |
| 14 | 94.5 | 92.39766081871345 | 90.3061224489796 | 93.26424870466322 |
| 17 | 93.0 | 92.39766081871345 | 90.3061224489796 | 90.67357512953367 |
| 19 | 91.0 | 91.2280701754386 | 86.22448979591837 | 88.60103626943005 |
| 21 | 89.0 | 89.47368421052632 | 83.6734693877551 | 88.60103626943005 |
| 24 | 87.0 | 86.54970760233918 | 79.08163265306122 | 88.08290155440415 |
| 26 | 86.5 | 84.7953216374269 | 77.0408163265306 | 86.01036269430051 |
| 28 | 85.0 | 83.04093567251462 | 71.42857142857143 | 82.90155440414507 |
| 31 | 76.5 | 78.94736842105263 | 58.6734693877551 | 72.02072538860104 |
| 33 | 73.0 | 76.60818713450293 | 54.59183673469388 | 69.94818652849742 |
| 35 | 70.5 | 74.26900584795322 | 50.0 | 64.24870466321244 |
| 38 | 61.0 | 70.76023391812865 | 37.244897959183675 | 53.8860103626943 |
| 40 | 58.0 | 64.32748538011697 | 31.632653061224488 | 47.668393782383426 |
| 42 | 54.0 | 59.64912280701755 | 21.42857142857143 | 39.37823834196891 |
| 45 | 47.0 | 49.70760233918129 | 17.346938775510196 | 29.53367875647669 |
| 47 | 38.0 | 44.44444444444444 | 13.775510204081627 | 25.388601036269435 |
| 49 | 31.0 | 42.10526315789473 | 11.734693877551024 | 18.13471502590673 |
| 51 | 24.5 | 31.578947368421055 | 9.183673469387756 | 14.507772020725383 |
| 53 | 18.5 | 26.31578947368422 | 6.122448979591837 | 9.844559585492235 |
| 55 | 14.0 | 19.88304093567251 | 2.040816326530617 | 4.663212435233163 |
| 57 | 4.5 | 12.280701754385973 | 0.0 | 2.5906735751295287 |
| 59 | 2.5 | 5.847953216374265 | 0.0 | 1.0362694300518172 |
| 61 | 2.0 | 3.5087719298245617 | 0.0 | 0.5181347150259086 |
| 63 | 1.0 | 1.1695906432748586 | 0.0 | 0.0 |C
D
E
F
#### Chart: Canton S Males + Histidine Lifespan
| Category | 0 mM | 1 mM | 5 mM | 10 mM |
|---|---|---|---|---|
| 0 | 100.0 | 100.0 | 100.0 | 100.0 |
| 2 | 100.0 | 99.49238578680203 | 100.0 | 99.5 |
| 4 | 100.0 | 98.98477157360406 | 100.0 | 99.0 |
| 6 | 99.5 | 97.96954314720813 | 100.0 | 98.5 |
| 8 | 99.5 | 97.46192893401015 | 100.0 | 98.0 |
| 10 | 99.0 | 97.46192893401015 | 98.5 | 98.0 |
| 12 | 99.0 | 96.95431472081218 | 98.0 | 97.5 |
| 14 | 98.5 | 96.44670050761421 | 97.0 | 96.0 |
| 17 | 98.0 | 95.93908629441624 | 96.5 | 94.0 |
| 19 | 96.5 | 95.93908629441624 | 96.0 | 94.0 |
| 21 | 96.0 | 92.89340101522842 | 95.0 | 94.0 |
| 24 | 96.0 | 91.37055837563452 | 92.0 | 90.0 |
| 26 | 95.0 | 89.84771573604061 | 89.5 | 88.0 |
| 28 | 92.0 | 87.30964467005076 | 86.5 | 85.0 |
| 31 | 86.0 | 80.71065989847716 | 82.5 | 80.5 |
| 33 | 83.5 | 74.11167512690355 | 77.5 | 72.0 |
| 35 | 80.0 | 70.55837563451777 | 71.5 | 65.5 |
| 38 | 67.5 | 64.46700507614213 | 63.5 | 57.5 |
| 40 | 58.5 | 55.83756345177665 | 54.5 | 51.0 |
| 42 | 52.5 | 47.71573604060914 | 49.0 | 43.50000000000001 |
| 45 | 37.5 | 39.086294416243646 | 40.5 | 33.0 |
| 47 | 26.5 | 32.99492385786802 | 32.5 | 26.0 |
| 49 | 18.5 | 26.39593908629442 | 26.0 | 16.5 |
| 51 | 14.0 | 20.812182741116743 | 18.0 | 9.5 |
| 53 | 8.0 | 13.705583756345177 | 10.5 | 5.5 |
| 55 | 6.0 | 5.583756345177662 | 4.0 | 2.0 |
| 57 | 3.5 | 0.507614213197968 | 2.0 | 0.0 |
| 59 | 3.5 | 0.0 | 1.0 | 0.0 |
| 61 | 1.5 | 0.0 | 0.5 | 0.0 |
| 63 | 0.0 | 0.0 | 0.0 | 0.0 |
#### Chart: w1118 Females + Histidine Lifespan
| Category | 0 mM | 1 mM | 5 mM | 10 mM |
|---|---|---|---|---|
| 0 | 100.0 | 100.0 | 100.0 | 100.0 |
| 2 | 98.3695652173913 | 99.45652173913044 | 98.53658536585365 | 100.0 |
| 4 | 94.56521739130434 | 97.82608695652173 | 97.5609756097561 | 98.99497487437186 |
| 6 | 92.93478260869566 | 97.28260869565217 | 94.63414634146342 | 98.49246231155779 |
| 8 | 92.93478260869566 | 95.65217391304348 | 94.63414634146342 | 97.98994974874373 |
| 10 | 91.30434782608695 | 92.3913043478261 | 91.70731707317073 | 95.47738693467336 |
| 12 | 91.30434782608695 | 91.84782608695652 | 90.73170731707317 | 94.9748743718593 |
| 14 | 89.67391304347827 | 91.84782608695652 | 88.78048780487805 | 93.96984924623115 |
| 17 | 88.04347826086956 | 89.67391304347827 | 86.82926829268293 | 91.95979899497488 |
| 19 | 86.95652173913044 | 89.67391304347827 | 84.8780487804878 | 87.93969849246231 |
| 21 | 82.6086956521739 | 86.95652173913044 | 82.4390243902439 | 83.41708542713567 |
| 24 | 77.17391304347827 | 86.41304347826087 | 79.51219512195122 | 76.88442211055276 |
| 26 | 72.28260869565217 | 82.6086956521739 | 77.07317073170731 | 73.36683417085428 |
| 28 | 66.30434782608695 | 76.63043478260869 | 72.6829268292683 | 67.33668341708542 |
| 31 | 57.608695652173914 | 55.97826086956522 | 61.46341463414634 | 59.79899497487437 |
| 33 | 53.26086956521739 | 52.17391304347826 | 47.3170731707317 | 54.77386934673367 |
| 35 | 46.7391304347826 | 48.91304347826087 | 37.5609756097561 | 52.26130653266331 |
| 38 | 36.41304347826087 | 33.69565217391305 | 22.92682926829268 | 45.7286432160804 |
| 40 | 29.347826086956516 | 27.173913043478265 | 17.5609756097561 | 39.69849246231156 |
| 42 | 24.456521739130437 | 21.73913043478261 | 12.682926829268297 | 27.1356783919598 |
| 45 | 16.304347826086953 | 1.6304347826086882 | 7.317073170731703 | 10.050251256281399 |
| 47 | 9.23913043478261 | 0.5434782608695627 | 2.439024390243901 | 6.030150753768851 |
| 49 | 9.23913043478261 | 0.0 | 0.48780487804877737 | 3.0150753768844254 |
| 51 | 5.978260869565219 | 0.0 | 0.0 | 1.5075376884422127 |
| 53 | 3.8043478260869534 | 0.0 | 0.0 | 0.5025125628140756 |
| 55 | 1.6304347826086882 | 0.0 | 0.0 | 0.0 |G
#### Chart: w1118 Males + Histidine Lifespan
| Category | 0 mM | 1 mM | 5 mM | 10 mM |
|---|---|---|---|---|
| 0 | 100.0 | 100.0 | 100.0 | 100.0 |
| 2 | 100.0 | 98.32402234636872 | 98.98989898989899 | 98.98477157360406 |
| 4 | 95.05494505494505 | 94.41340782122904 | 94.44444444444444 | 94.9238578680203 |
| 6 | 94.50549450549451 | 92.17877094972067 | 93.43434343434343 | 93.90862944162437 |
| 8 | 93.4065934065934 | 89.94413407821229 | 90.4040404040404 | 93.40101522842639 |
| 10 | 91.75824175824175 | 88.26815642458101 | 88.38383838383838 | 91.87817258883248 |
| 12 | 90.65934065934066 | 86.59217877094972 | 87.87878787878788 | 91.37055837563452 |
| 14 | 89.01098901098901 | 84.91620111731844 | 86.86868686868686 | 89.84771573604061 |
| 17 | 86.81318681318682 | 81.56424581005587 | 84.84848484848484 | 86.29441624365482 |
| 19 | 84.06593406593407 | 78.2122905027933 | 83.83838383838383 | 83.248730964467 |
| 21 | 82.41758241758242 | 76.53631284916202 | 78.78787878787878 | 81.21827411167513 |
| 24 | 73.62637362637362 | 68.71508379888269 | 67.17171717171718 | 66.497461928934 |
| 26 | 70.87912087912088 | 62.56983240223464 | 57.57575757575758 | 60.91370558375635 |
| 28 | 66.4835164835165 | 58.100558659217874 | 52.525252525252526 | 56.85279187817259 |
| 31 | 37.362637362637365 | 45.81005586592178 | 37.37373737373737 | 40.10152284263959 |
| 33 | 33.51648351648352 | 41.340782122905026 | 34.848484848484844 | 36.54822335025381 |
| 35 | 28.57142857142857 | 37.43016759776536 | 31.313131313131322 | 32.99492385786802 |
| 38 | 13.736263736263737 | 18.994413407821227 | 14.141414141414145 | 14.720812182741113 |
| 40 | 12.637362637362642 | 13.40782122905027 | 12.121212121212125 | 12.182741116751274 |
| 42 | 10.439560439560438 | 8.938547486033528 | 8.080808080808083 | 9.137055837563452 |
| 45 | 2.7472527472527446 | 1.6759776536312927 | 3.030303030303031 | 0.507614213197968 |
| 47 | 1.098901098901095 | 1.1173184357541857 | 2.525252525252526 | 0.0 |
| 49 | 1.098901098901095 | 1.1173184357541857 | 2.525252525252526 | 0.0 |
| 51 | 1.098901098901095 | 0.558659217877107 | 0.0 | 0.0 |
| 53 | 1.098901098901095 | 0.0 | 0.0 | 0.0 |
| 55 | 0.0 | 0.0 | 0.0 | 0.0 |
| 57 | 0.0 | 0.0 | 0.0 | 0.0 |

### Slide 11
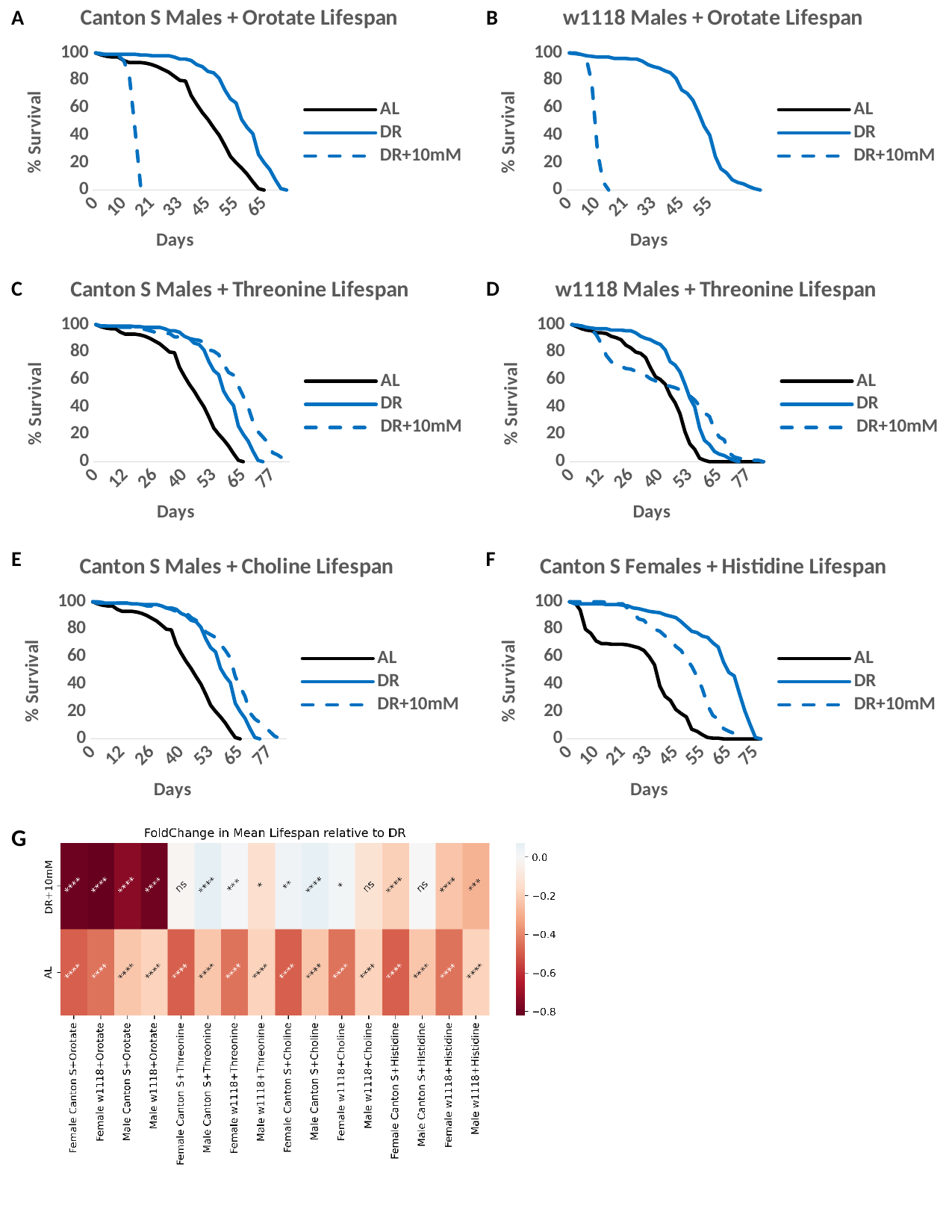

[unsupported chart]
[unsupported chart]
A
B
[unsupported chart]
[unsupported chart]
C
D
E
[unsupported chart]
F
[unsupported chart]
G
